# Yanomami skin microbiome complexity challenges prevailing concepts of healthy skin

**DOI:** 10.1101/2024.11.23.625008

**Authors:** Juliana Durack, Yvette Piceno, Hoang Vuong, Brian Fanelli, David A. Good, Nur A. Hasan, Manoj Dadlani, Larry Weiss, Julia Oh, Aleksandar D. Kostic, Thomas L Dawson, Hortensia Caballero-Arias, Rita R. Colwell

## Abstract

The adult skin microbiome contributes to skin homeostasis and generally comprises relatively low microbial complexity, especially sebaceous sites, where lipophilic *Cutibacterium* and *Malassezia spp.* predominate. Current understanding of the healthy skin microbiome derives predominantly from studies of western, industrialized populations, with limited representation of diverse cultures and lifestyles. In this study, the skin microbiome of a remote indigenous Yanomami community was investigated, revealing a complex microbial community comprised of 115 novel bacterial genomes. The bacterial community composition included genera common to western skin microbiota and additional diverse taxa, which formed multiplex interactions with a dominant eukaryote, *Malassezia globosa*. Metatranscriptome-derived functional attributes of the microbial communities contributed to skin homeostasis, fortifying barrier integrity via lipid metabolism and acid production and protecting against oxidative stress. The Yanomami skin microbiome comprised a suite of microbial taxa co-detected within their various surroundings. Longitudinal monitoring of Western expeditioner microbiome revealed acquisition of the Yanomami skin microbiome following immersion into the Amazon and subsequent loss upon return to an industrialized setting. These findings challenge the prevailing Western-centric view of what comprises a healthy adult skin microbiome, suggesting a diverse community that includes bacteria of environmental origin confers benefits not recognized in the current model of healthy skin. Importantly, we highlight the malleability of adult skin microbiome composition, which includes changes sustained by lifestyle modification.

## Introduction

The human skin serves as a protective external barrier and primary communication with the environment. Skin microbiota shape and maintain multiple physiological processes in this outermost layer of the human body^1^. The role of the skin microbiome includes maintenance of an acidic pH that protects against pathogen colonization^2^, establishing and modulating immune tolerance^3–6^, and fortifying skin barrier integrity^7–9^. Site-specific microenvironments are established by local pH, temperature, moisture, sebum content, topography, and other factors that influence community structure. Sebaceous (oily) surfaces are dominated by lipophilic microorganisms, specifically *Cutibacterium spp* and *Malassezia spp*, and differ from those found at more diverse and less stable dry or moist skin sites^10–12^. Site-specific interpersonal variation of skin microbiomes has also been linked with various host traits and habits, including age, sex, BMI, cohabitation, smoking, pet ownership, and antibiotic use^12–19^. Environment and lifestyle, specifically urbanization, also influence the skin microbiota, exemplified by the observation that individuals residing in rural communities harbor a more diverse and compositionally distinct microbiome than their urban-dweller counterparts^18,20–23^.

Perturbations to cutaneous microbial communities have been linked to various skin conditions, such as acne, rosacea, psoriasis, and atopic dermatitis^24–26^. These pathologies are considered industrialized comorbidities with less frequent occurrence in developing nations and unreported in hunter-gatherer populations ^27–29^. The frequency of cutaneous disease inversely correlates with cutaneous microbial diversity observed along an urbanization gradient^22,23^. The least impacted populations are those of seminomadic hunter-gatherers with the highest skin microbial diversity^22,23,30–32^. Since the current understanding of the role of the skin microbiome in human health derives predominantly from studies of Western, industrialized populations, it remains unclear if assumptions generated in Western-biased populations apply across all cultures and lifestyles.

The Yanomami are amongst the last remaining swidden horticulturalists, hunter-gatherers indigenous people of the Amazon who maintain minimal contact with non-Yanomami outsiders. This study focused on understanding the composition and functional attributes of the Yanomami skin microbiome of a remote community that remains unaltered by industrialization.

### Study Population and Design

The Yanomami are people indigenous to the Amazon rainforests of Venezuela and Brazil, traditionally living a swidden horticulturalist, seminomadic hunter-gatherer lifestyle. Samples for this study were collected in one such community in Venezuela from two neighboring shabono (communal living structure – comprised of a naturally derived thatched roof and an open-aired shelter, with no dividing walls and a natural forest ground as the floor), situated within walking distance of a river where community members fish, bathe and collect water for drinking and cooking. To respect the Yanomami decision-making procedure that is followed for participation in the study, a community meeting was held where the field team explained each stage of sample collection, the goal, and the scope of the microbiome research project. To ensure an intercultural perspective, didactic posters and pictures were used with all the information translated into the Yanomami language. Community concurrence and free and prior informed verbal consent were obtained, after which sampling commenced.

A total of 94 skin samples were collected for metagenomic analysis from 17 Yanomami (50% female; 1-70 approximate years of age). During two expeditions, individuals from the same community were sampled at seven skin sites: retroauricular crease (RAC), face, upper back, scalp, volar arm, axilla, and toe web space (Table S1), with no significant difference between expeditions in coverage of age groups or body sites sampled (Chi-square p=0.319 and p=0.443, respectively). The same body sites of a Western adult male expeditioner were longitudinally sampled at various time points throughout the journey from the metropolitan United States to a remote Yanomami community and return, yielding a total of 45 skin samples (Table S2). To compare indigenous skin microbiomes with those of Western industrialized individuals, skin samples from the expeditioner (excluding those collected while in the Amazon) and 16 RAC samples (44% female; age range 18-40yo) from the Human Microbiome Project (HMP)^33^ data archive were included in the analysis. Median de-hosted metagenomic read depth was 2.17E7, 2.52E7 reads/sample for Yanomami skin samples and HMP samples, respectively. As proxy for sebaceous skin microbiome transcriptional activity, RNA extracted from chest swabs collected from Yanomami adults and Westernized expeditioner (Table S2) was profiled for microbial composition and functional analysis. To elucidate the relationship of Yanomami skin microbiota composition to the surrounding environment, samples of soil from an entrance trail to the shabono, water samples from a creek and river, and ash from the heath located in the shabono were collected and included in the study. Lastly, a Yanomami adult female (Yanomami traveler) accompanied the expeditioner on his return to the USA, and samples from her RAC, scalp, and volar arm were collected approximately two weeks into an extended stay in an industrialized setting, providing contrast with samples collected while in the native Amazon community (Table S2, for study schema, see Fig S1).

## Results

### Skin metagenomes of the Yanomami harbor novel microbial genomes

We first assessed percentage of human reads across combined samples from the Yanomami RAC and face (n=30), which were significantly lower (ANOVA of linear mixed-effects model (LME) p<0.001) compared to the expeditioner westernized samples (USA samples, n=10) collected from the same sebaceous body sites (Fig 1A). This trend persisted across all body sites, although statistical significance was reached only for RAC, scalp, and arm (Table S3), indicating possible reduced desquamation and potentially stronger skin barrier. Surprisingly, a lower percentage of bacterial (but not fungal) reads was detected in the Yanomami skin samples, resulting in a higher percentage of remaining unmapped reads (ANOVA of LME p<0.001; Fig 1A and Table S3). This observation was consistent when the data were normalized to a read depth of 13M, after removing human reads (Fig S2A). A portion of unmapped reads from Yanomami RAC/face samples presented as novel bacterial species, akin to observations of gut microbiomes reported for other hunter-gatherers^34^. Through reference-independent de novo assembly, reconstruction of bacterial metagenome-assembled genomes (MAGs) recovered 115 novel ( ≥50% completion and ≤5% contamination) bacterial genomes, constituting 59.0% of the total reconstructed prokaryotic MAGs obtained from the samples (Fig 1B; Table S4) when classified ( ≥90% average nucleotide identity) using The Genome Taxonomy Database (GTDB) ^35–37^ or 155 novel (79.5%) MAGs (FigS2B) when classified using The Skin Microbial Genome Collection (SMGC)^38^. The largest number of novel genomes were identified as *Corynebacterium* (12.3% and 13.3% of all MAGs, using GTDB or SMGC, respectively), with several genomes identified only to family (or higher), such as *Dermatophilaceae*, *Micrococcaceae* and *Propionibacteriaceae*. Additionally, 18 (≥50% BUSCO completion, dereplicated) fungal MAGs were recovered from the same samples, most of which were identified as *Malassezia*, including two potentially novel strains identified as *Malasseziaceae* (Table S5). These results suggest that the Yanomami skin harbors an assortment of unique microbes yet to be identified and characterized.

**Figure 1:**
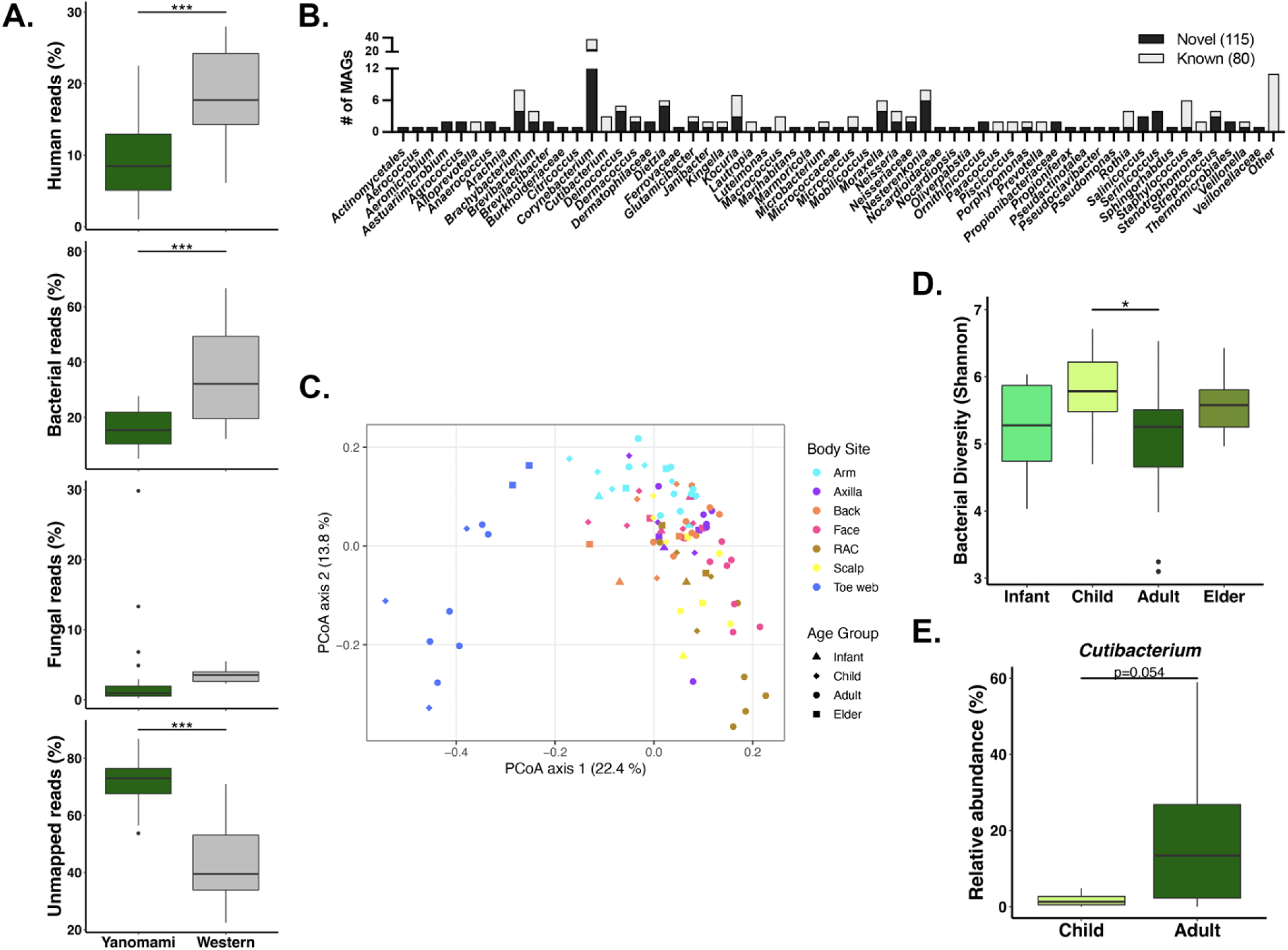
**A.** Percent of raw read distribution in RAC and face samples in Yanomami (n=30) and Western expeditioner (n=10; ANOVA of LME p-value: ***<0.001). **B.** GTDB-based classification of bacterial MAGs reconstructed from Yanomami RAC/face samples. **C.** Principal Coordinates Analysis (Bray-Curtis) of Yanomami (n=94) bacterial microbiota across body sites and age groups. **D.** Bacterial Diversity (Shannon Index) of combined sebaceous samples (RAC/face/back/scalp; n=53) across age groups (Tukey multiple comparisons: * p<0.05). **E.** Relative abundance of *Cutibacterium* in combined sebaceous samples (n=40) in Yanomami children vs adults. (ANOVA of LME)

### The Yanomami skin microbiome harbors diverse bacterial and fungal communities

We next assessed compositional and functional differences between Yanomami and Western skin microbiomes. The microbial populations were analyzed by mapping to known microbial genomes using normalized 13M reads after de-hosting. Similar to industrialized skin microbiomes, factors such as body site^10–12^ and age ^13–16,19^ explained a portion of the variation in bacterial (Nested PERMANOVA estimated component of variation (ECV)=20.4%, p=0.0001 and ECV=9.4%, p=0.004, respectively Table S6 and Fig1C) and fungal (Nested PERMANOVA ECV=20.9%, p=0.0001 and ECV=10.1%, p=0.024; Table S6 and FigS2C) Yanomami skin composition, where communities clustered primarily by microenvironment. The bacterial diversity of sebaceous skin (back, face, RAC and scalp) showed age-related variance, mirroring observations reported for industrialized skin. Specifically, bacterial diversity declined in adulthood (Fig 1D, Tukey multiple comparisons p<0.05), based on increased relative abundance of *Cutibacterium spp.* (Fig 1E, ANOVA of LME p=0.054; FigS2D and FigS3A). Fungal diversity was also lower in adult skin microbiota compared to that of children (FigS3B, ANOVA of LME p<0.05), corresponding to a higher relative abundance of *Malassezia spp.*, (FigS3C-D, ANOVA of LME p<0.05). The decline in diversity in adult skin was not reflected in the richness of bacterial or fungal communities on combined sebaceous sites (data not shown), nor was it significant for dry skin (arm) samples from the same subjects (FigS3E-F), indicating the community signature is primarily associated with restructuring rather than membership loss.

Compared to the Western skin microbiome, bacterial communities of the Yanomami adult RAC were significantly richer (80% [IC range 76-87%]), more diverse (67% [IC range 60-74%], Wilcoxon, p=<0.001, Fig2A), and compositionally distinct (PERMANOVA R2=0.455, p=0.001, Fig2B), with less dominance by *Cutibacterium spp.* (Fig2C). Higher bacterial diversity on Yanomami skin was not exclusive to RAC; it was also evident on other sebaceous body sites when contrasted with the Westernized expeditioner samples (FigS4A-C), as well as on the arm (dry skin site; FigS4D). Bacterial diversity did not differ significantly in samples from the toe web (moist body site; FigS4E), although composition remained distinct. The most abundant bacterial taxa detected on Yanomami skin were species of *Corynebacterium* (*C. mucifaciens* and *C. ureicelerivorans*), *Bordetella* (*B. pertussis*), *Kocuria* (*K. indica* and *K. palustris*), *Micrococcus* (*M. aloeverae* and *M. luteus*), *Brachybacterium* (*B. paraconglomeratum*), *Microbacterium,* and *Moraxella* (*M. osloensis*; Fig2C-D). At the genus level, a wide range of bacterial taxa were absent or only infrequently detected on Western skin. These included species of *Brachybacterium, Brevibacterium, Microbacterium,* and *Janibacter*, amongst others (Fig2E), that were detected on Yanomami skin irrespective of the age of subjects. A subset of genera was frequently detected in both groups, including species of *Corynebacterium* and *Cutibacterium* (Fig2E), although in variable relative abundance (FigS5A). The relative abundance of *Staphylococcus*, another shared genus, was similar for both groups (Wilcoxon with BH-correction q>0.05). The repertoire of *Staphylococcus spp.* was greater on Yanomami skin, with significantly higher relative abundance of *S. aureus*, *S. hominis*, *S. haemolyticus, S. saprophyticus,* and *S. arlettae* (Fig2D), but no significant difference for *S. epidermidis* (FigS5A). Notable, virulence factors of the *Staphylococcus spp.,* expressed as a ratio of virulence genes to the relative abundance of species within the genus, trended lower for Yanomami (FigS5B). Yanomami *Staphylococcus* virulence factors involved attachment and colonization, such as the surface protein SdrC-E (FigS5C), whereas virulence factors more common on Western skin included antibiotic resistance and pathogenesis, namely ThyA and SePI (FigS5C).

**Figure 2:**
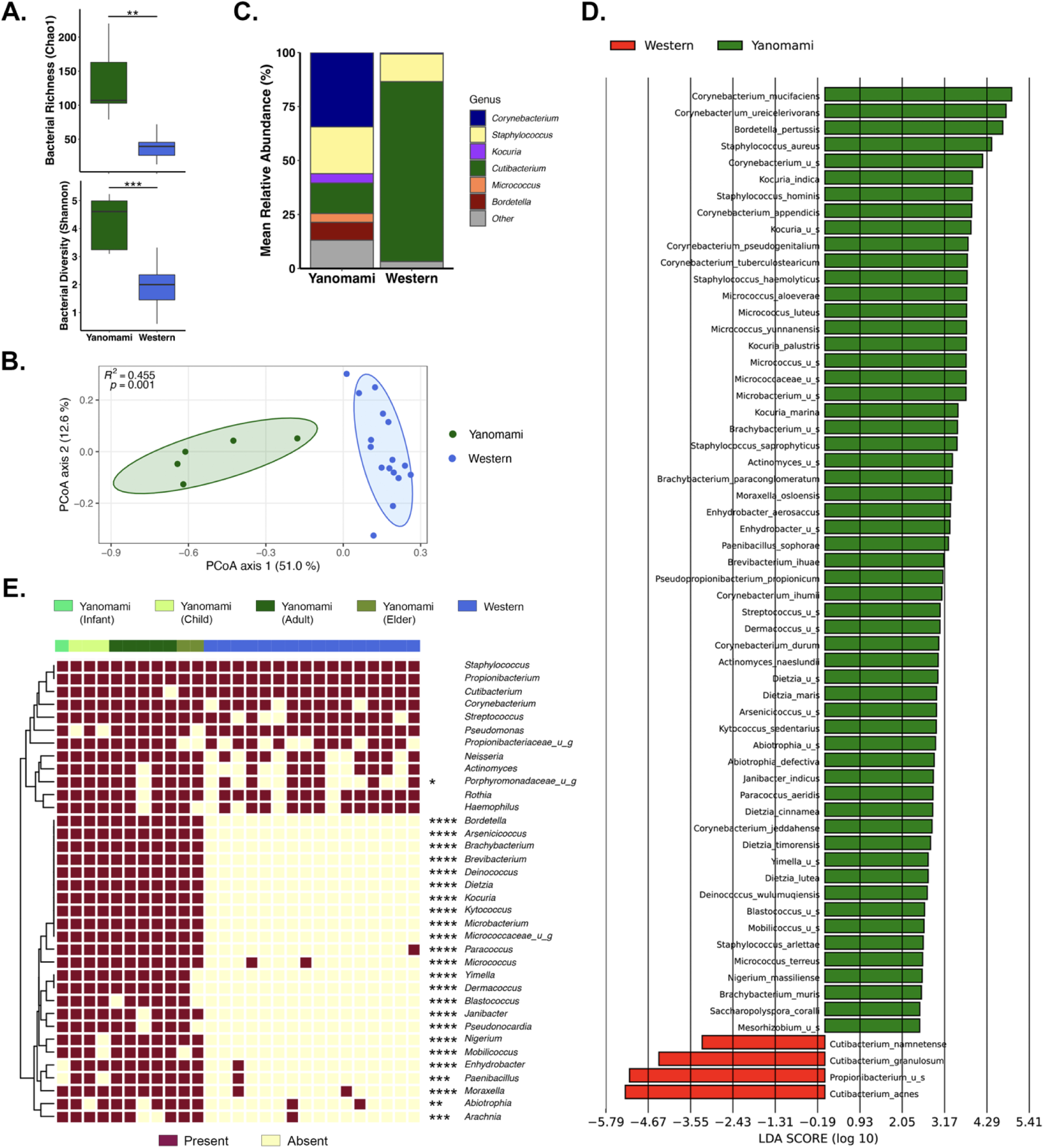
Bacterial community composition of Yanomami adult RAC is divergent from westerners. **A.** Alpha Diversity of Yanomami (n=5) and HMP (n=16; Wilcoxon test p-value: ***<0.001, **<0.01). **B.** Beta diversity (PERMANOVA, Bray-Curtis). **C.** Mean relative abundance of dominant (≥2%) genera. **D.** Most differential species (LEfSe, at >75% of subjects per group). **E.** Distribution of genera present at >80% (Yanomami n=11; color indicates age group; Fisher’s exact test, FDR-adjusted: **** q<0.0001, *** q<0.001, ** q<0.01, * q<0.05).

Relative abundance of bacteriophages reflected the bacterial profile (FigS5D), with *Propionibacterium* phages significantly more abundant on Western skin (Wilcoxon with Benjamini-Hochberg correction q<0.05). The relative abundance of *Propionibacterium* phage correlated with the relative abundance of *Cutibacterium* (formerly *Propionibacterium*) on Western (Spearman correlation, rho=0.47, p=0.016) but not Yanomami (rho=-0.14, p>0.1) skin. A greater variety of *Propionibacterium* phages was detected on Western skin (FigS5E), although the Yanomami skin microbiome harbored a higher relative abundance of the *Propionibacterium* phage PFR2 (FigS5E).

Fungal communities of both groups were dominated by *Malassezia spp.* (Fig3A). However, the Yanomami mycobiota of the adult RAC samples were richer (Wilcoxon, p=<0.01, FigS6A) and compositionally distinct (PERMANOVA R^2^=0.376, p=0.004, FigS6B-C), but trended less diverse due to dominance by *M. globosa* (Fig3A-B and FigS6D). Trending richer fungal communities were detected on all Yanomami sebaceous sites sampled (FigS7A-C), though fungal diversity trends for those sites were mixed compared to that of the Western expeditioner. No significant difference in alpha diversity matrices was observed for the arm (dry site; FigS7D) and toe web mycobiota (moist site; FigS7E), although all body sites were compositionally distinct, with reduced abundance of *M. restricta* (FigS6D; S7A-E and Fig3A). Other fungal species differentiating Yanomami RAC samples included *M. japonica*, *M. furfur* and *Brettanomyces bruxellensis* (Fig3B and FigS6C).

**Figure 3:**
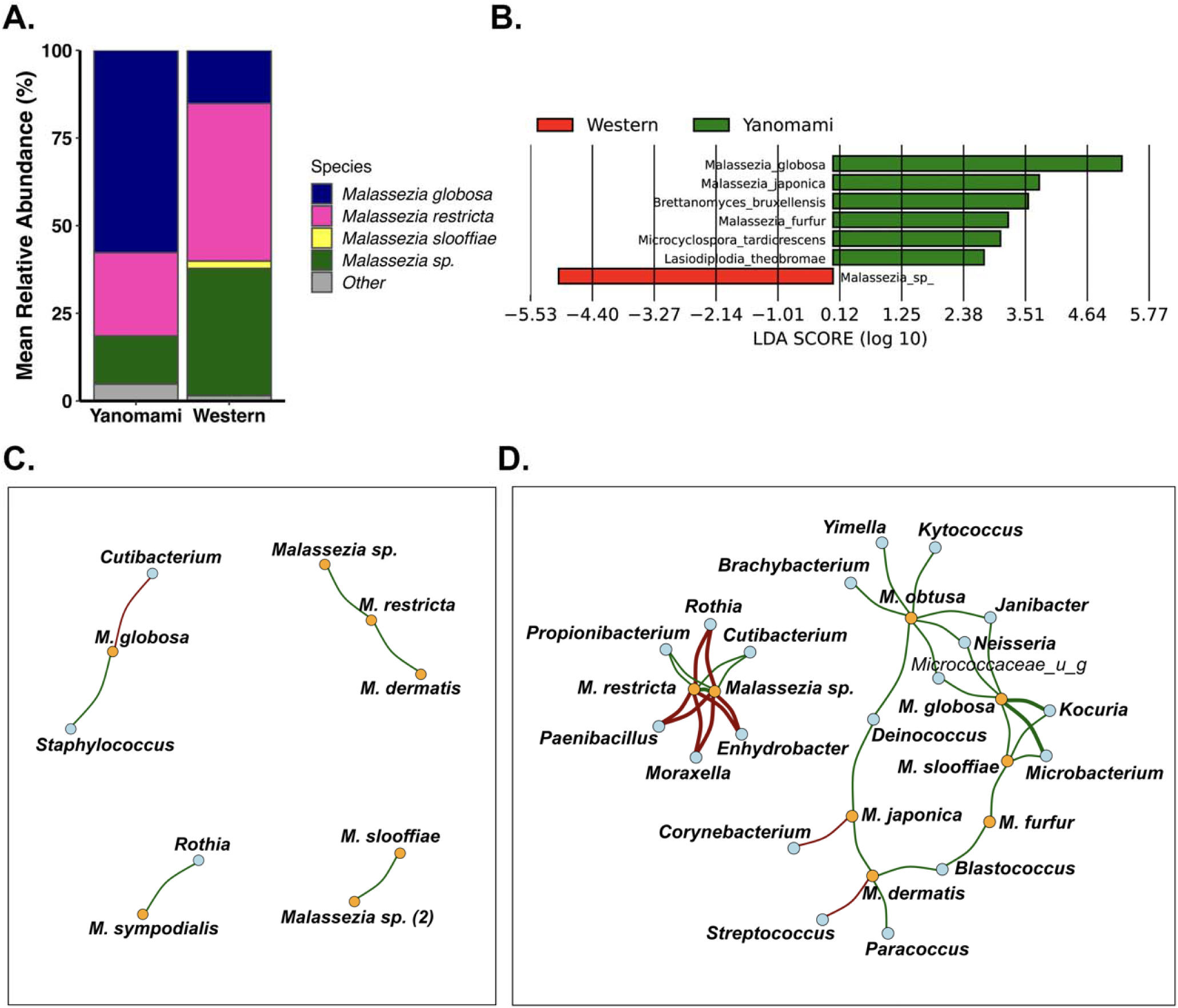
Fungal community composition of Yanomami adult RAC is divergent from westerners and comprises distinct interkingdom interactome. **A.** Mean relative abundance of dominant species (≥2%) in Yanomami (n=5) vs HMP (n=16) individuals. **B.** Most differential taxa (LEfSe;>50% of subjects/group). Co-occurrence of dominant bacterial genera (80% prevalence) and all *Malassezia* species in the **C.** western HMP and **D.** Yanomami skin microbiomes (Spearman [abundance score], p<0.05; where: Red - negative interactions; Green - positive interactions; thickness of lines indicates co-association strength).

### The Yanomami skin microbiome features distinct interkingdom interactions and functional attributes

The interkingdom dynamics of the Yanomami skin microbiome were compared to Western skin using network analysis of abundance scores, interactions of bacterial genera with members of the most dominant fungi, *Malassezia,* and interspecies interactions within this genus. The adult Yanomami RAC skin microbiome exhibited a more complex interactome than Western skin. Whereas *M. globosa* was directly co-associated with *Cutibacterium* (negatively) and *Staphylococcus* (positively) on Western skin (Fig3C), on Yanomami skin, *M. globosa* was co-associated with several bacterial genera, including *Kocuria, Microbacterium, Neisseria,* and *Janibacter* (Fig3D). Additionally, *M. globosa* was associated with *M. slooffiae* and indirectly with *M.obtusa*, forming a highly interactive hub (Fig3D). Conversely, *M. restricta* was observed to inversely correlate with *Moraxella*, *Enhydrobacter, Paenibacillus*, *Rothia* and directly with *Cutibacterium* (Fig3D). Relative abundance of *M. globosa* correlated positively with overall bacterial richness and diversity on the skin irrespective of lifestyle (FigS8A-C), the inverse having been observed for *M. restricta*.

Reflective of compositional differences, the functional potential of the Yanomami skin microbiome was both greater than (Fig4A) and distinct from the Western microbiome (PERMANOVA R^2^=0.47, p=0.001, Fig4B). Functional pathways of the Yanomami skin microbiome that were most differential included lipid metabolism (biosynthesis and degradation), fermentation, metabolite activation, degradation of various amides/amines, and hormone metabolism (Fig4C). In contrast, Western skin microbiome pathways included cofactor biosynthesis and degradation of carbohydrates and amino acids (Fig4C). To determine if these functional attributes were expressed on the skin, the Yanomami skin microbial transcriptome was compared to that of the Western expeditioner, based on an additional sebaceous site (chest, exclusively utilized for RNA extraction and transcriptomics). The Yanomami transcriptional profile revealed richer, more diverse (FigS9A) and compositionally distinct bacterial (FigS9B-C) and fungal (FigS9D-F) communities, with comparable membership (FigS9C and FigS9F) as observed from the metagenomic analyses, suggesting metabolically active microbes on Yanomami skin.

**Figure 4:**
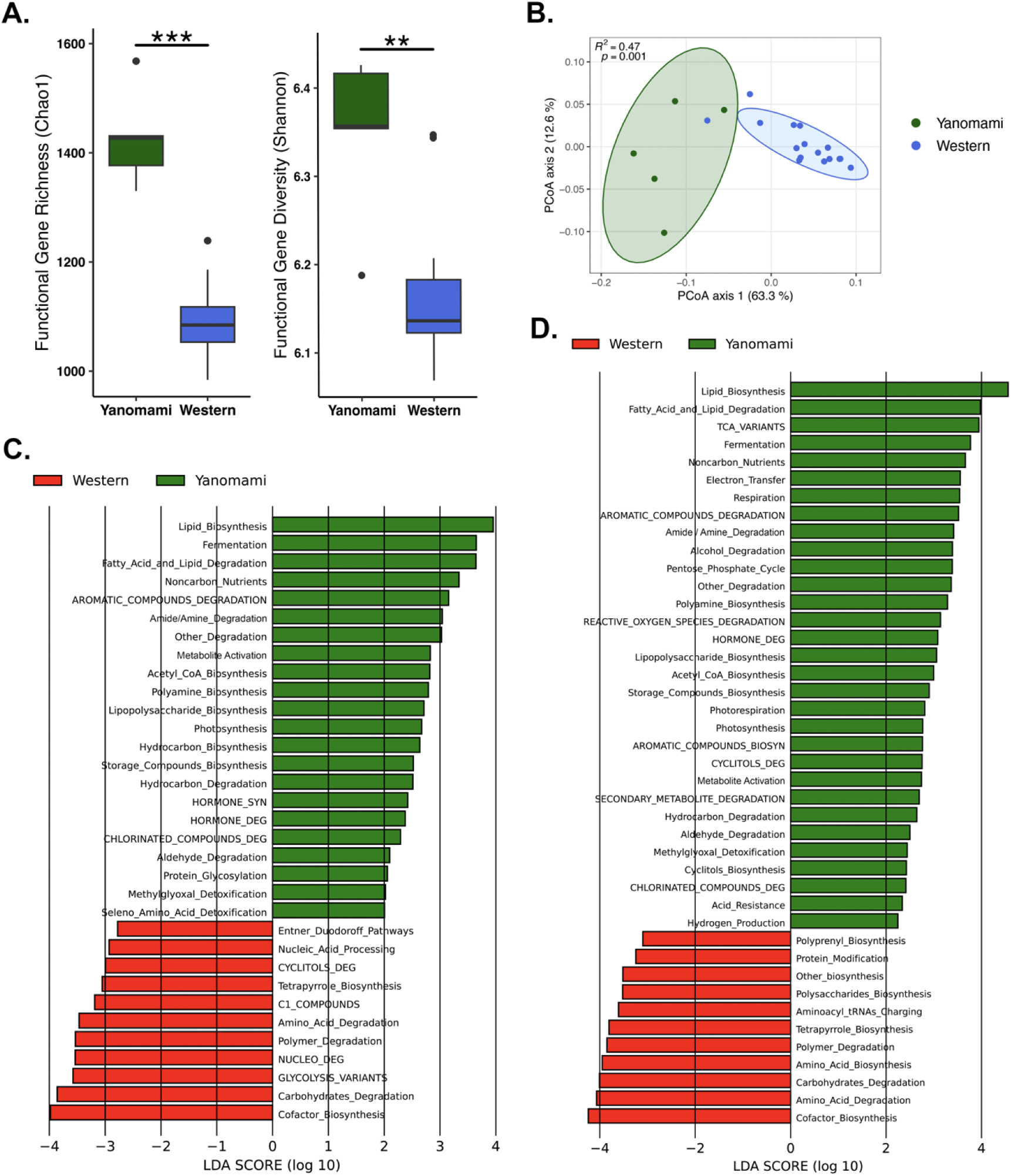
Functional attributes of the Yanomami adult microbiome are distinct from western individuals **A.** Richness (obs) and diversity (Shannon index) of functional potential (3512 pathways assigned by MetaCyc) of the Yanomami (RAC, n=5) and western (RAC, n=16) skin microbiome. Wilcoxon test p-value: ***<0.001, **<0.01. **B.** Functional potential composition (PERMANOVA of Bray-Curtis dissimilarity; pathways assigned by MetaCyc) of RAC samples. **C.** Most differential functional secondary pathway classes in RAC samples (LEfSe). **D.** Most differential functional secondary pathway classes of active microbiota in chest (RNA) samples of Yanomami adults (n=4) and western expeditioner (n=3) samples (LEfSe).

Transcriptomics-based functional community output of the Yanomami microbiome was more complex and functionally divergent (FigS9G-H) from that of the Western microbiome. Active functional pathways of the Yanomami microbiome most differential were lipid biosynthesis, fatty acid and lipid degradation, fermentation, aromatic compound metabolism, degradation of amides/amines, hormone, and reactive oxygen species (Fig4D). Western skin microbial functional pathways included cofactor biosynthesis, amino acid metabolism, and carbohydrate degradation (Fig4D), suggesting physiologically divergent states of functional activity in these compositionally distinct microbial communities.

### The Yanomami skin microbiome is integrated with their environment and diminished by industrialization

The Yanomami skin microbiome varied in bacterial and fungal composition (FigS10A-H) across body sites (arm, axilla, back, face, RAC, and toe web), but unlike Western skin ^10–12^, greatest bacterial community richness was detected in the Yanomami face (sebaceous site) samples and the least in toe web (moist site, FigS10A-B) samples, the latter also harboring the least rich fungal communities (FigS10D-E). Arm (dry site) and axilla (moist site) microbiota shared significant community composition with sebaceous skin samples (FigS10C and FigS10F). Observations likely related to Yanomami’s lifestyle and the close connection to the environment.

The Yanomami live in communal shabono open to the environment and regularly swim in and use local water sources. Microbial exchange with their surroundings is likely a contributor to the microbial richness of the Yanomami skin microbiome. Yanomami skin microbiota across all body sites, clustered more closely with that of soil that had been sampled from an entrance trail to the shabono rather than that of the creek (Bray-Curtis distance matrix; FigS11B), with toe web microbiota being most compositionally similar to that of soil. The more abundant cutaneous bacterial genera, namely *Pseudomonas* and *Staphylococcus,* were detected in both environmental profiles (Fig5A), *Brachybacterium*, *Deinococcus*, *Janibacter*, *Microbacterium* and *Micrococcus* as well as others (Fig5A; FigS11A) were detected in soil. The mycobiome was reflective of creek microbial communities (FigS11C-D) except toe web, unsurprisingly most similar in composition to soil, with *M. restricta* and *Brettanomyces bruxellensis* in creek and soil samples, respectively.

**Figure 5:**
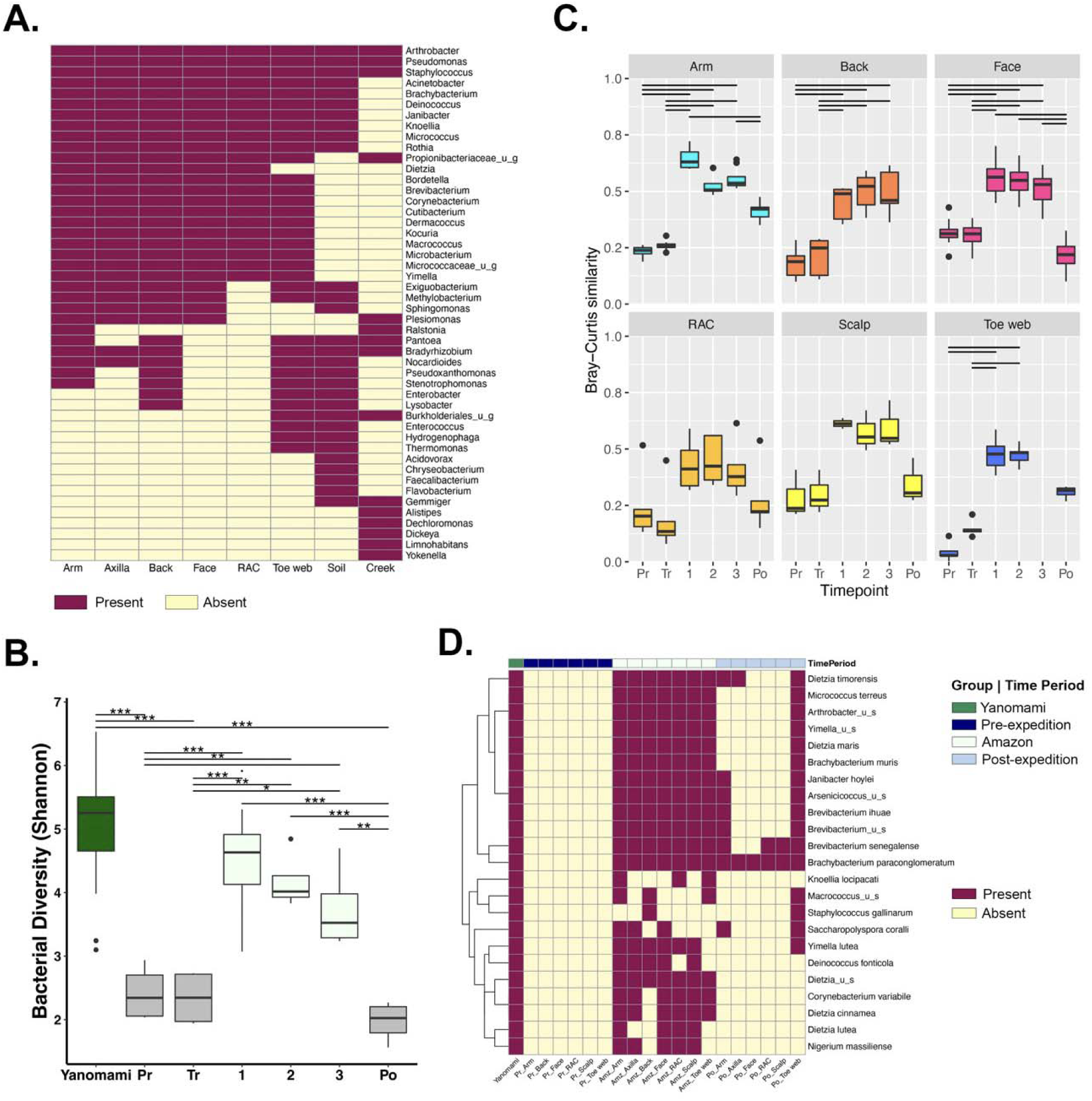
The Yanamami skin microbiome is interconnected with their environment and acquirable upon adoption of local practices**. A.** Distribution of bacterial genera (≥2%, in at least three adult Yanomami) at various body sites that are co-detected in or exclusive to environmental samples. **B.** Comparison of bacterial diversity of adult Yanomami (n=9, combined sebaceous sites, n=26 samples) and western expeditioner (n=23 same body sites) at various timepoints; Pr-pre-expedition, Tr-in-transit, 1-Amazon, 2-Amazon, 3-Amazon, Po-post-expedition; ANOVA of LME p=0.02, (Tukey: *** p<0.001; ** p<0.01; * p<0.05). **C.** Similarity in bacterial composition (Bray-Curtis) between the western expeditioner and adult Yanomami (Friedman p<0.05, Dunn test [pair-wise, BH-corrected p<0.025]). **D.** Bacterial species present in ≥25% Yanomami adults at ≥0.25% relative abundance co-detected on the expeditioner at various time points.

Further evidence of environmental influence on the Yanomami skin microbiota was observed from the longitudinally sampled skin microbiome of the expeditioner, who adopted the local practices in the daily activities of the Yanomami community. An increase in microbial complexity was noted in samples from combined sebaceous body sites at all three time points of collection in the Amazon (approximately one week apart), with bacterial richness (FigS11E), diversity (Fig5B), and composition closely resembling that of Yanomami microbiota (Fig5C-D). Apparent fungal richness gain for the Amazon samples did not reach statistical significance (FigS11F), likely due to *Malassezia* dominance. The Yanomami cutaneous microbial community complexity diminished after integration into industrialized lifestyle, both in the microbiota of the expeditioner (Fig5D and FigS11E-F) and a Yanomami family member who traveled out of the Amazon (FigS12A-C), and was accompanied by loss of several bacterial genera namely *Dietzia*, *Kocuria*, *Micrococcus*, *Brevibacterium*, *Brachybacterium*, *Deinococcus*, *Yimella* and *Janibacter*, all prominent members of the Yanomami cutaneous microbiota (Fig5D and FigS12C).

## Discussion

The Yanomami represent one of the few indigenous groups in the Amazon remaining minimally impacted by industrialization, leading a seminomadic, swidden-horticulturalist, hunter-gatherer lifestyle. They experience few chronic cutaneous inflammatory conditions ^27–29^, in contrast to an increasing frequency in industrialized populations globally. The Yanomami skin microbiome has been described (based on 16S rRNA biomarker sequencing) as significantly diverse compared to industrialized microbiome^22,23,30,32^, but inadequately characterized. This study aimed to elucidate the composition and functional attributes of the Yanomami skin metagenome and determine whether Western-biased assumptions about the skin microbiome apply across cultures and lifestyles, especially as drastically different as the Yanomami, relatively unaffected by industrialization.

The Yanomami sebaceous skin metagenome was found to be complex, with a large component of total reads not mappable to human, fungal, bacterial, or viral genomes in the current literature and available databases, likely reflecting a connection to the diverse and relatively unexplored rainforest biome^39^ where the Yanomami reside. The Yanomami skin harbored a unique assortment of microbes with more than half of the reconstructed bacterial metagenome-assembled genomes (MAGs) belonging to previously unidentified microorganisms, including those co-evolved with mammalian and human skin^40^, classified only to family level, such as *Dermatophilaceae*, *Micrococcaceae*, *Propionibacteriaceae and Malasseziaceae*. The skin microbiome is reported as co-evolving in response to environmental and host cues, where microbial assemblages vary by skin site and are conditioned by the local cutaneous microenvironment, a relationship observed in the study reported here for Yanomami skin. Like that of industrialized skin^10–16,19^, the Yanomami skin bacterial and fungal community composition is influenced by site microenvironment and age of participants, with a bloom and gradual decrease in lipophilic *Cutibacterium* and *Malassezia* relative abundance with age. Community restructuring was reflected in a significant decline in microbial diversity in adults but not in bacterial richness and fungal assemblies on sebaceous sites, nor on dry skin for the same subjects, perhaps related to age-associated hormonal changes and increased sebum production in adulthood^16^.

Unlike industrialized sebaceous skin of relatively low microbial complexity due to dominance by *Cutibacterium* and *Malassezia*, where skin health correlates with less complex microbial communities^41^, the Yanomami adult skin microbiome was significantly more diverse. It was enriched for a wide range of environmental bacteria absent or infrequent on Western skin, such as *Brachybacterium, Brevibacterium, Microbacterium,* and *Janibacter spp*. A subset of bacterial genera was common to both populations, such as *Cutibacterium, Staphylococcus,* and *Corynebacterium spp.,* with the latter most abundant on Yanomami skin and representing novel MAGs. *Corynebacterium spp.* are key immunomodulatory members of the cutaneous microbiome^42^ and have previously been reported to support community diversity and evenness in species distribution within the modern skin microbiome^43^.

*Staphylococcus* was detected in both Western and indigenous adult skin microbiomes. Although its relative abundance was indistinguishable, species diversity was greater on Yanomami skin, with a higher relative abundance of *S. aureus*, *S. hominis*, *S. haemolyticus*, *S. gallinarum,* and *S. saprophyticus*, with diminished virulence potential compared to Western *Staphylococcus*. While Yanomami *Staphylococcus* were putatively enriched for factors related to attachment and colonization, virulence attributes of Western *Staphylococcus* included factors related to antibiotic resistance and enhanced pathogenesis, such as *S. aureus* thymidylate synthase^44^ and *S. epidermidis* pathogenicity islands^45^. Bacteriophage populations in this study proved unremarkable, possibly restricted by available bacteriophage sequences in public databases primarily derived from clinically relevant bacteria. Reflective of the bacterial profile, *Propionibacterium* phages were more abundant on the Western skin, except for lytic phage, PFR2, previously isolated from *P. freudenreichii,* with reported ability to infect acne-associated *C. acnes*^46^ as a species abundant on the Yanomami skin.

The sebaceous adult skin mycobiota was dominated by the lipophilic yeast *Malassezia*. However, *M. globosa* and *M. restricta* dominated the skin mycobiome of the Yanomami and Western skin, respectively. *M. globosa* was negatively co-associated with *Cutibacterium* in the Western skin samples (as previously reported^47^) and positively with *Staphylococcus*. On the Yanomami skin, *M. globosa* formed a positive interaction hub, directly co-associating with several bacterial genera. The relative abundance of *M. globosa* correlated positively with overall bacterial richness and diversity on the skin of Yanomami and Western subjects. Collectively, these observations indicate that *M. globosa* relative abundance may be a marker of greater bacterial complexity in sebaceous skin microbiota and, perhaps, as a gatekeeper of skin health. There is growing evidence that *M. globosa* may be beneficial to healthy skin in preventing colonization by pathogenic microorganisms^48–50^ and modulating inflammation via induction of immunoregulatory cytokine IL-10^51^. It may have a protective role in dermatitis, as suggested by the clinical observation of a lower *M.restricta/globosa* ratio on healthy skin compared to subjects with seborrheic dermatitis and dandruff, two conditions where *Malassezia* is implicated^52^.

Microbial diversity and interspecies interactions contribute to the functional output of microbial communities. Hence, it was unsurprising to observe a higher functional potential and more transcriptionally active microbial assemblages on Yanomami than Western skin. Of the functional pathways that most differentiated the Yanomami microbiomes were those with the potential to contribute to maintaining skin homeostasis and fortifying skin barrier integrity. These included lipid metabolism and fermentation, contributing to pH and antimicrobial defensins regulation on the skin surface^53,54^. Degradation of secreted aromatic compounds and hormones by environmental bacteria^55^ abundant on the Yanomami skin may diffuse microbial virulence of pathogenic species^56^, whilst enhanced physiologic REDOX function renders additional protection against oxidative stress, such as that induced by UV exposure^57,58^.

Our findings suggest that similar to industrialized skin, where bacterial interconnections are observed with the indoor environment^59,60^ in which, on average, 90% of westerners spend their days^61^, the Yanomami skin microbiome is interconnected with their natural environment, with a suite of key differential cutaneous bacterial taxa co-detected across various environmental samples. Furthermore, longitudinal sampling of a Western expeditioner skin microbiota throughout a journey to a remote Yanomami community provided evidence that the microbial complexity of the Yanomami skin may be acquired upon adoption of local practices. Temporary establishment of bacteria from contact with soil and aquatic environments has been reported on Western skin^62–66^. Here, we conclude that prolonged and continuous exposure will result in incorporation of environmental taxa into an established skin microbiome. Our observation of the loss of the Yanomami skin bacterial complexity following adoption of a Western lifestyle agrees with the report of bacterial diversity loss along an urbanization gradient in a cross-sectional assessment of the skin microbiome in the same geographic region^22,23^.

In summary, results of this study underscore lifestyle as a primary determinant in shaping the composition of cutaneous microbiota, with industrialization likely a pivotal factor driving the loss of environmental microbes such as those that enrich the Yanomami skin microbiome. The complex microbial communities observed in this study to comprise the indigenous skin microbiome challenge current Western-centric and biased views of a healthy adult skin microbiome, whereby skin health is deemed correlating with less complex microbial communities. The findings reported here support the hypothesis that a diverse skin microbiota enriched with environmental microbes may confer benefits not captured by the current model of healthy skin. Finally, we provide evidence suggesting that the adult human skin microbiome may be stably altered by prolonged microbial exposure, giving hope that continuous topical microbial supplementation can stabilize and substantially improve skin health by modulating the skin microbiome.

## Limitations of the study

While the study represents the most comprehensive analysis of non-industrialized skin microbiome survey to date, it is limited to a relatively small sample size, a single Yanomami community and a Western expeditioner, all of which was constrained by logistics, strict conformation to ethical practices for study of the last remaining minimally contacted hunter-gatherer population in the Amazonian rainforest, and lack of cutaneous metagenome studies of other populations maintaining a similar lifestyle for comparative reference. Other limitations of the study included the large number of uncharacterized genes, two-fold greater in Yanomami metagenomes and the lack of closely related homologs in currently available public databases, restricting putative functional prediction of genes in these communities. To address these limitations, studies involving deeply sequenced skin metagenomics of underrepresented non-westernized populations are needed for a more comprehensive understanding of the diversity and functional attributes of the human skin microbiome.

## Methods

### Inclusion & Ethics Statement

All samples used in this study were collected legally under valid permits from the Venezuelan government and ethically by The Yanomami Foundation (formally The Good Project) a nonprofit organization and was reviewed and approved by the Institutional Board of Directors, Venezuelan Institute of Scientific Research (IVIC; DIR-0021/1582/2017) and Advarra’s (formally IntegReview) Institutional Review Board (IORG0000689). Permission to collect samples was obtained from community leaders and each adult or parent of a child by verbal consent with the help of a translator and by video recording when possible. No written consent was acquired since this Yanomami community does not have a written form of communication. To minimize disturbance to the Yanomami way of life, sample collection was performed by a single Yanomami researcher, who was integrated into community’s daily activities. Western expeditioner provided a written informed consent.

This study was made possible by the dedication of local researchers at IVIC who enabled the acquisition of bioethical and legal permits for conducting research in the Yanomami territory of Venezuela and recruitment of the Yanomami participants in the study, as well as facilitated transport of the microbiome samples from Venezuela to the USA. David Good is a member of a Yanomami tribe and the executive director of the Yanomami Foundation, he personally collected the samples used in this study. The Yanomami Foundation is the custodian of the Yanomami Microbiome Project. Their mission is to equip the Yanomami communities with the tools, resources, and intercultural training necessary to safeguard the Yanomami way of life and preserve their Amazonian lifestyle and their microbiome. The Foundation does this partly by facilitating and advancing international, transdisciplinary collaborative research of the Yanomami microbiomes, believing this elucidates links between the unique Yanomami microbiome and their health and well-being. Importantly, this illuminates the vital importance of preserving the Amazonian rainforest and the intimate connection the Yanomami have with it. The Foundation sets a new precedent in bioethics by using community-based participatory approaches, exchanging intercultural skills and training, and ensuring equitable benefit sharing with the Yanomami communities in Venezuela and Brazil (for more detail of how the foundation supports local communities, visit www.yanomamifoundation.org).

### Sample Collection

Cutaneous microbiome samples were collected from arm, axilla, back, chest, face, retroauricular crease (RAC), scalp, and toe web using swabs (Isohelix, Kent, UK) and preserved in DNA/RNA Shield (Zymo Research, Irvine, CA). Environmental samples were collected using swabs (hearth ash, soil, water) or Sterivex filters (water) and preserved in DNA/RNA Shield (Zymo Research, Irvine, CA). Sample collection was performed by a single individual following IRB-approved protocols (detailed in Supplemental Information).

### Nucleic acid extraction and metagenomic sequencing

DNA was extracted from back swabs using ZymoBIOMICs MagBead DNA Kit (Zymo Research, Irvine, CA, USA), and ZymoBIOMICS Microprep kit (with Metapolyzyme enzymatic pre-treatment) was used for the other body site sample collection. Qiagen PowerSoil Pro kit (Qiagen, Germantown, MD, USA) was used to extract DNA from environmental samples. RNA was extracted from chest swabs and environmental samples (river, soil, and ash) using Qiagen RNeasy Plus kit (Qiagen, Germantown, MD, USA). DNA sequencing library was prepared using Nextera XT library prep kit (Illumina, San Diego, CA, USA), as previously described^67^. RNA sequencing library was prepared using the SMARTer Stranded RNA Total RNA Sample Prep Kit - Pico Input (Takara Bio USA, San Jose, CA, USA), as previously described^68^. All sequencing was done using 2 x 150bp using Illumina HiSeq 4000 Instrument (Illumina Inc., San Diego, CA, USA).

### Data analysis

Unassembled metagenomic sequencing reads from the dataset and HMP westerners were analyzed using CosmosID bioinformatics software package (CosmosID Inc., Rockville, MD, United States), as described previously^69^. Human reads were removed using genome assembly GRCh38_p6 and normalized by subsampling to 13M reads using *reformat.sh* of the BBTools suite (http://bbtools.jgi.doe.gov). Processed reads were classified to taxa using CosmosID portal, employing k-mer mapping to unique genome regions for each taxon^67,69^ (details in Supplemental Information). Reads were assigned to bacterial and archaeal, fungal, protist, viral (excluding phage), or bacteriophage lineages for taxonomic comparisons. Positive control taxa were removed from the dataset prior to analysis. Possible process contaminants were identified (Table S7) as skin and environment in origin but not removed from the dataset. For functional gene classification, quality-filtered reads were translated and used to search the UniRef90 database^70^. Gene families were annotated using MetaCyc^71^, and normalized abundance values were calculated as copies per million (see Supplemental Information for details).

### Metagenome assembly and taxonomic assignment

Metagenome-assembled genomes (MAGs) from face and RAC samples were assembled with ‘metaWRAP’^72^. Adapters and host reads were removed, reads were assembled with metaSPAdes^73^ for prokaryotic MAGs, the remaining unassembled reads were assembled with MEGAHIT^74^, and contigs were aligned (SAMtools^75^ and BWA-MEM^76^) and binned (MetaBAT2^77^, MaxBin 2.0^78^, and CONCOCT^79^; see details in Supplemental Information). Following bin refinement, bacterial representative genomes (n=195, with ≥50% completion and ≤5% contamination) were dereplicated (dRep) ^80^. and classified with GTDB-tk^81^ (v2.10 07-RS207) as known (≥95% average nucleotide identity [ANI] to a genome from the GTDB^36,37^) or novel. The same ANI criteria were applied to bacterial MAGs compared against the Skin Microbial Genome Collection (SMGC)^38^.

Fungal contigs assembled with metaSPAdes^73^ were retrieved using EukRep (west et al 2018), indexed and aligned (SAMtools^75^ and BWA-MEM^76^), and binned (MetaBAT2^77^). Fungal representative genomes (n= 18; ≥50% BUSCO^82^ completion) were dereplicated (dRep)^80^ and classified with taxator-tk^83^ and Blast+^84^ as known (≥95% ANI of NCBI assembly) or novel.

### Statistical analyses

Univariate data comparisons between Yanomami and Western individuals were performed using Wilcoxon rank test, Fisher’s Exact Test, or Analysis of variance (ANOVA) of linear mixed-effects (LME) models^85^ (lmerTerst package in R) to account for repeated measures, where appropriate. Univariate comparisons among Yanomami age groups were performed using Kruskal-Wallis, followed by Dunn test for multiple pairwise comparisons. Multiple comparisons p-values were adjusted using Benjamini-Hochberg^86^ (BH) correction (q-values). Multivariate data comparisons were performed using permutational analysis of variance (PERMANOVA^87^, *adonis* in R) on Bray-Curtis (BC) dissimilarity matrices (square root-transformed taxonomic data) and visualized using Principal Coordinates Analysis (PCoA). Linear discriminant analysis Effect Size^88^ (LEfSe) algorithm provided between-group comparisons of taxa or functional gene relative abundances. Spearman correlation (*Hmisc* in R) of abundance scores was used to identify multi-kingdom co-occurrences. *Malassezia* species interactions to bacterial genera were plotted using *igraph* in R when p<0.05, as previously described^89^. Differences across adult Yanomami body sites and similarities between environmental samples and adult Yanomami skin microbiota were evaluated by the Friedman test (group-wise comparison) and the Dunn test. Longitudinal evaluation of the western expeditioner’s microbiota at combined sebaceous sites relative to the Yanomami was performed using LME models, followed by Tukey pairwise comparisons. Details for all analyses and analysis software used are provided in the supplementary information.

## Data Availability

Raw data from metagenomic and metatranscriptomic analyses generated in this study have been deposited in the NCBI under BioProject number PRJNA1083578.

## Acknowledgements

We are indebted to the Yanomami people for entrusting the stewardship of the Yanomami Microbiome Project to the Yanomami Foundation who enabled this study. The authors wish to acknowledge Monica Contreras (Instituto Venezolano de Investigaciones Científicas, IVIC) for support in the recruitment of Yanomami participants and navigating the permit process in Venezuela. Roger Cariban, coordinator of the Indigenous Health of Amazonas State (Ministerio del Poder Popular para la Salud, MPPS) for his medical oversite and assistance with sample collection. Orlana Lander (Instituto de Medicina Tropical, Universidad Central de Venezuela) for assistance with sample collection. Chris Callewaert (Ghent University) and Nayeim Khan (CosmosID) for assistance with exploratory analysis. As well as Martin Feelisch (University of Southampton), Howard Maibach (University of California, San Francisco), Emma Allen-Vercoe (University of Guelph), Carlos Bustamante (Stanford University; GalateaBio) for useful data discussions.

## Author Contributions

JD, YP, LW contributed to study conception and design. DAG and HCA managed bioethics and legal permit acquisition and served as cultural liaisons for the Yanomami people. DAG, HCA, JD, YP contributed to sample collection and management. JD, YP, HV, BF, NAH, MD, JO, ADK, TLD contributed to data acquisition, analysis and interpretation; YP, HV, BF, JD performed data analysis. All authors have contributed to manuscript drafting and have approved the final version of the manuscript.

