## Supplemental Tables and Figures for "Yanomami skin microbiome complexity challenges prevailing concepts of healthy skin"

**Table S1:** Yanomami cohort demographic and sample collection distribution.

|  |  | **Samples collected** | | | | | | | |
| --- | --- | --- | --- | --- | --- | --- | --- | --- | --- |
| **Demographic Characteristics** | **Individuals** | **RAC** | **Face** | **Back** | **Scalp** | **Arm** | **Axilla** | **Toe web** | **Chest (RNA)** |
| **Total No.** | **18^*^** | **12** | **18** | **14** | **9** | **18** | **12** | **11** | **12** |
| ***Age Group^#^*** |  |  |  |  |  |  |  |  |  |
| Infant (<2) | **2** | 1 | 2 | 1 | 1 | 2 | 1 | 0 | 2 |
| Child (2-12) | **5** | 3 | 5 | 3 | 3 | 5 | 3 | 3 | 4 |
| Adult (16-50) | **9^*^** | 6^*^ | 9^*^ | 8^*^ | 3^*^ | 9^*^ | 6^*^ | 6^*^ | 4 |
| Elder (>50) | **2** | 2 | 2 | 2 | 2 | 2 | 2 | 2 | 2 |
| ***Sex*** |  |  |  |  |  |  |  |  |  |
| Male | **9** [4]^^^ | 5 [3]^^^ | 9 [4]^^^ | 5 [3]^^^ | 3 [2]^^^ | 9 [4]^^^ | 5 [3]^^^ | 4 [3]^^^ | 7 [2] ^^^ |
| Female | **9** [5]^^^ | 7 [3]^^^ | 9 [5]^^^ | 9 [5]^^^ | 6 [1]^^^ | 9 [5]^^^ | 7 [3]^^^ | 7 [3]^^^ | 5 [2] ^^^ |
| ***Expedition^Ψ^*** |  |  |  |  |  |  |  |  |  |
| Expedition1 | **12** [4]^^^ | 6 [1]^^^ | 12 [4]^^^ | 8 [3]^^^ | 8 [3]^^^ | 12 [4]^^^ | 6 [1]^^^ | 5 [1]^^^ | 12 [4] ^^^ |
| Expedition2 | **6** [5]^^^ | 6 [5]^^^ | 6 [5]^^^ | 6 [5]^^^ | 1 [0]^^^ | 6 [5]^^^ | 6 [5]^^^ | 6 [5]^^^ | 0 |

^#^Groups were assigned based on approximate age (years). ^*^One individual adult male was sampled in both expeditions. ^^^numbers in square brackets refer to break down of samples collected from adult individuals. ^Ψ^No significant difference in the coverage of age groups nor body site between two expeditions (Chi-square p=0.319 and p=0.443, respectively).

**Table S2:** Sample distribution for the Western expeditioner and the Yanomami traveler.

|  | **Samples collected** | | | | | | | |
| --- | --- | --- | --- | --- | --- | --- | --- | --- |
| **Western expeditioner** | **RAC** | **Face** | **Back** | **Scalp** | **Arm** | **Axilla** | **Toe web** | **Chest (RNA)** |
| Pre-expedition | 1 | 1 | 2 | 1 | 1 | 1 | 1 | 1 |
| In transit^*^ | 3 | 2 | 3 | 2 | 4 | 1 | 2 | 1 |
| Yanomami Village^#^ | 5 | 3 | 3 | 3 | 3 | 3 | 2 | 0 |
| Post-expedition | 1 | 2 | 0 | 1 | 1 | 1 | 1 | 1 |
| *Expedition:* |  |  |  |  |  |  |  |  |
| Expedition1 | 8 | 7 | 6 | 6 | 6 | 5 | 5 | 3 |
| Expedition2**^^^** | 2 | 1 | 2 | 1 | 3 | 1 | 1 | 0 |
| **Yanomami Traveler** |  |  |  |  |  |  |  |  |
| Yanomami Village | 1 | 1 | 1 | 1 | 1 | 1 | 1 | 1 |
| Westernized | 1 | 0 | 0 | 1 | 1 | 0 | 0 | 0 |

^*^Samples were collected pre- and post- Amazon in urban settings outside of metropolitan US. **^#^**Sample collection was conducted at three time points throughout the duration of the stay. **^^^**Only collected samples in transit (prior to or after leaving the Amazon) during this expedition.

**Table S3** - Distribution of Human, Bacterial, Fungal, and unmapped reads for Yanomami vs western skin at each of the sampled body sites.

|  |  |  | **% Human** | | |
| --- | --- | --- | --- | --- | --- |
| **Body Site** | **#y** | **#w^&^** | **y** | **w^&^** | **p-value^*^** |
| RAC | 12 | 5 | **17.9** [11.4-26.1] | **42.2** [32.0-65.8] | **0.006** |
| Face | 18 | 5 | **21.8** [14.5-36.76] | **35.9** [19.7-58.3] | 0.199 |
| Back | 14 | 4 | **15.9** [6.3-30.7] | **37.8** [11.5-82.1] | 0.442 |
| Scalp | 9 | 5 | **6.5** [4.1-19.5] | **69.1** [42.9-82.1] | **0.004** |
| Arm | 18 | 6 | **5.6** [4.0-8.7] | **28.9** [17.1-37.8] | **0.004** |
| Axilla | 12 | 4 | **5.7** [4.0-8.9] | **25.8** [3.0-65.1] | 0.684 |
| Toe Web | 11 | 4 | **2.1** [1.5-3.3] | **6.8** [3.3-10.4] | *0.056* |
|  |  |  | **% Bacterial** | | |
| **Body Site** | **#y** | **#w^&^** | **y** | **w^&^** | **p-value^*^** |
| RAC | 12 | 5 | **17.0** [14.3-24.5] | **30.7** [15.8-42.1] | 0.130 |
| Face | 18 | 5 | **13.1** [8.8-19.7] | **45.7** [18.6-62.5] | **0.009** |
| Back | 14 | 4 | **9.1** [7.1-15.9] | **39.0** [20.6-70.7] | **0.005** |
| Scalp | 9 | 5 | **12.3** [10.5-15.9] | **7.5** [4.3-27.2] | 0.519 |
| Arm | 18 | 6 | **12.7** [8.7-19.8] | **21.4** [18.8-29.9] | **0.015** |
| Axilla | 12 | 4 | **21.5** [11.9-26.0] | **19.7** [6.2-39.7] | 1.000 |
| Toe Web | 11 | 4 | **2.6** [1.5-8.7] | **34.0** [24.3-36.4] | **0.002** |
|  |  |  | **%Fungal** | | |
| **Body Site** | **#y** | **#w^&^** | **y** | **w^&^** | **p-value^*^** |
| RAC | 12 | 5 | **1.7** [0.5-4.4] | **3.6** [3.1-4.8] | 0.124 |
| Face | 18 | 5 | **0.8** [0.3-1.4] | **2.9** [2.4-3.8] | **0.001** |
| Back | 14 | 4 | **0.4** [0.3-0.6] | **1.6** [0.8-2.4] | **0.017** |
| Scalp | 9 | 5 | **1.3** [0.4-3.0] | **5.7** [3.1-11.6] | **0.012** |
| Arm | 18 | 6 | **0.3** [0.2-0.5] | **12.6** [7.4-4.1] | **0.000** |
| Axilla | 12 | 4 | **0.3** [0.2-0.8] | **0.7** [0.3-3.3] | 0.303 |
| Toe Web | 11 | 4 | **0.1** [0.08-0.1] | **0.1** [0.04-0.4] | 0.951 |
|  |  |  | **%Unmapped** | | |
| **Body Site** | **#y** | **#w^&^** | **y** | **w^&^** | **p-value^*^** |
| RAC | 12 | 5 | **70.8** [63.4-75.9] | **41.6** [35.3-62.3] | **0.009** |
| Face | 18 | 5 | **73.4** [68.6-77.3] | **36.4** [23.5-52.7] | **0.000** |
| Back | 14 | 4 | **82.3** [75.3-83.8] | **36.4** [21.4-45.8] | **0.001** |
| Scalp | 9 | 5 | **79.7** [72.1-83.9] | **51.3** [43.6-57.8] | **0.001** |
| Arm | 18 | 6 | **80.9** [71.6-85.7] | **46.2** [40.5-47.8] | **0.000** |
| Axilla | 12 | 4 | **73.9** [68.3-82.3] | **54.3** [37.7-64.0] | **0.009** |
| Toe Web | 11 | 4 | **95.4** [89.7-97.5] | **61.5** [57.3-72.6] | **0.002** |

**^&^** Westernized samples collected from expeditioner were included as western comparison group. **^*^**Significance was determined using Mann-Whitney test.

**Table S4:** Summary of bacterial metagenome-assembled genomes (MAGs) reconstructed from RAC/Face samples and classified using The Genome Taxonomy Database.

| **Taxonomy Group^&^** | **Class^#^** | **Median Completeness** | **Median Contamination** | **Median**  **Strain Heterogeneity** | **# of Dereplicated MAGs in Group** | **MedianGenome Size** | **MedianN50** | **Median Contig #** |
| --- | --- | --- | --- | --- | --- | --- | --- | --- |
| Actinomyces | K | 79.07 | 0.91 | 0 | 1 | 2347886 | 12980 | 320 |
| Pseudactinotalea | N | 73.05 | 1.36 | 33 | 1 | 3163011 | 7734 | 532 |
| Bifidobacterium | K | 100.00 | 0.09 | 0 | 1 | 2203580 | 119991 | 41 |
| Brevibacterium | K | 94.55 | 1.74 | 100 | 1 | 2561230 | 11399 | 301 |
| Brachybacterium | K | 82.70 | 1.24 | 12 | 4 | 2881743 | 7881 | 481 |
| Brachybacterium | N | 84.07 | 0.66 | 20 | 4 | 2471961 | 11283 | 323 |
| Dermatophilaceae | N | 71.31 | 0.91 | 17 | 2 | 2501380 | 8324 | 852 |
| Dermacoccus | K | 91.44 | 1.62 | 0 | 1 | 2640843 | 17103 | 279 |
| Dermacoccus | N | 91.57 | 0.52 | 50 | 2 | 2821626 | 37493 | 181 |
| Janibacter | K | 88.95 | 0.54 | 100 | 1 | 2780532 | 19944 | 202 |
| Janibacter | N | 85.81 | 0.18 | 100 | 1 | 2911778 | 16240 | 299 |
| Kytococcus | K | 78.54 | 0.00 | 0 | 1 | 1963830 | 15571 | 274 |
| Marihabitans | N | 89.95 | 2.02 | 44 | 1 | 2440525 | 11758 | 333 |
| Ornithinicoccus | N | 85.26 | 0.97 | 58 | 2 | 3081740 | 18511 | 261 |
| Piscicoccus | K | 81.77 | 0.68 | 67 | 2 | 3804124 | 7781 | 807 |
| Serinicoccus | N | 71.21 | 0.62 | 0 | 4 | 2603655 | 7519 | 440 |
| Yimella | K | 95.60 | 1.08 | 50 | 1 | 2787536 | 28409 | 128 |
| Actinomycetales | N | 94.19 | 0.09 | 0 | 1 | 2871146 | 23999 | 209 |
| Agrococcus | N | 60.51 | 1.99 | 69 | 2 | 1667298 | 4113 | 460 |
| Microbacterium | K | 89.57 | 1.94 | 20 | 1 | 2419365 | 8939 | 348 |
| Microbacterium | N | 76.57 | 1.17 | 25 | 1 | 1888328 | 6194 | 385 |
| Pseudoclavibacter | N | 82.76 | 0.10 | 0 | 1 | 1954385 | 6400 | 401 |
| Micrococcaceae | N | 78.85 | 1.22 | 20 | 1 | 2237102 | 17978 | 145 |
| Citricoccus | N | 69.28 | 0.00 | 0 | 1 | 2085142 | 6279 | 411 |
| Glutamicibacter | K | 65.90 | 1.95 | 14 | 1 | 2436844 | 7307 | 405 |
| Glutamicibacter | N | 76.08 | 1.49 | 8 | 2 | 2476145 | 12794 | 313 |
| Kocuria | K | 95.08 | 0.00 | 0 | 4 | 2504947 | 27585 | 165 |
| Kocuria | N | 86.77 | 1.97 | 70 | 3 | 2511399 | 28522 | 130 |
| Micrococcus | K | 76.10 | 0.90 | 14 | 2 | 2097827 | 6917 | 468 |
| Micrococcus | N | 86.81 | 0.92 | 33 | 1 | 2060382 | 9553 | 317 |
| Nesterenkonia | K | 91.05 | 1.10 | 25 | 2 | 2356965 | 21832 | 185 |
| Nesterenkonia | N | 89.24 | 1.34 | 42 | 6 | 2287835 | 13807 | 237 |
| Rothia | K | 98.06 | 0.11 | 0 | 3 | 2432709 | 177352 | 125 |
| Rothia | N | 83.22 | 0.91 | 20 | 1 | 1812064 | 10475 | 246 |
| Corynebacterium | K | 89.69 | 0.62 | 33 | 13 | 2264999 | 15842 | 180 |
| Corynebacterium | N | 84.57 | 0.84 | 29 | 24 | 1958344 | 20810 | 149 |
| Dietzia | K | 64.29 | 1.76 | 50 | 1 | 2236012 | 13837 | 219 |
| Dietzia | N | 78.62 | 0.10 | 0 | 5 | 2313598 | 16099 | 232 |
| Lawsonella | K | 95.00 | 0.00 | 0 | 1 | 1655578 | 92883 | 43 |
| Nocardioidaceae | N | 87.63 | 0.95 | 0 | 1 | 2638100 | 21198 | 211 |
| Aeromicrobium | N | 88.62 | 0.52 | 50 | 1 | 2360823 | 32327 | 189 |
| Marmoricola | N | 89.35 | 1.28 | 0 | 1 | 2869920 | 14088 | 295 |
| Propionibacteriaceae | N | 79.58 | 0.13 | 50 | 2 | 2655552 | 17896 | 201 |
| Aestuariimicrobium | N | 84.50 | 1.75 | 75 | 2 | 2524088 | 12856 | 319 |
| Arachnia | N | 58.04 | 0.52 | 0 | 1 | 1581355 | 4049 | 440 |
| Brevilactibacter | N | 77.48 | 1.06 | 17 | 4 | 2443419 | 8357 | 409 |
| Cutibacterium | K | 95.54 | 0.12 | 0 | 3 | 2104395 | 45475 | 87 |
| Propioniferax | N | 89.77 | 2.94 | 11 | 1 | 2626599 | 8465 | 422 |
| Nocardiopsis | N | 70.60 | 0.18 | 0 | 1 | 4693416 | 4259 | 1269 |
| Alloprevotella | K | 76.38 | 0.98 | 30 | 2 | 1942301 | 14520 | 373 |
| Prevotella | K | 82.19 | 0.25 | 0 | 2 | 2350835 | 11121 | 344 |
| Porphyromonas | K | 97.53 | 0.00 | 0 | 1 | 2317207 | 85790 | 88 |
| Porphyromonas | N | 56.92 | 0.31 | 0 | 1 | 1130306 | 2580 | 472 |
| Thermomicrobiales | N | 70.82 | 0.57 | 83 | 2 | 2013707 | 4183 | 641 |
| Deinococcus | K | 66.03 | 2.00 | 33 | 1 | 2231124 | 3267 | 842 |
| Deinococcus | N | 64.37 | 0.67 | 42 | 4 | 2372150 | 4613 | 565 |
| Agathobacter | K | 98.73 | 1.53 | 14 | 1 | 2819909 | 9021 | 494 |
| Oliverpabstia | N | 57.02 | 3.51 | 0 | 1 | 1654495 | 5735 | 349 |
| Gemmiger | K | 98.64 | 0.00 | 0 | 1 | 2781203 | 55917 | 118 |
| Eubacterium | K | 52.68 | 0.95 | 50 | 1 | 815588 | 2820 | 620 |
| Anaerococcus | N | 75.23 | 2.09 | 6 | 2 | 1263640 | 6634 | 265 |
| Veillonellaceae | N | 83.08 | 0.22 | 0 | 1 | 1255874 | 13874 | 127 |
| Veillonella | K | 79.28 | 0.30 | 50 | 1 | 1288709 | 7362 | 232 |
| Veillonella | N | 96.41 | 0.13 | 0 | 1 | 1774152 | 13597 | 234 |
| Abiotrophia | K | 55.08 | 1.38 | 40 | 1 | 1128332 | 2658 | 906 |
| Aerococcus | N | 95.79 | 1.10 | 0 | 1 | 1787507 | 42647 | 54 |
| Lactiplantibacillus | K | 71.62 | 0.31 | 0 | 1 | 2403359 | 9331 | 316 |
| Streptococcus | K | 60.97 | 0.83 | 64 | 1 | 1241639 | 2944 | 522 |
| Streptococcus | N | 86.29 | 2.43 | 14 | 3 | 1628650 | 6502 | 228 |
| Salinicoccus | N | 80.83 | 0.99 | 0 | 3 | 1893507 | 6264 | 425 |
| Macrococcus | K | 78.66 | 1.10 | 60 | 3 | 1530300 | 4673 | 425 |
| Staphylococcus | K | 67.76 | 1.25 | 34 | 5 | 1750152 | 19565 | 223 |
| Staphylococcus | N | 53.79 | 3.45 | 0 | 1 | 1377389 | 2631 | 531 |
| Paracoccus | K | 67.98 | 0.81 | 17 | 2 | 2145028 | 4161 | 616 |
| Sphingorhabdus | N | 54.95 | 1.16 | 33 | 1 | 1546696 | 2924 | 585 |
| Burkholderiaceae | N | 85.35 | 0.41 | 0 | 1 | 3560623 | 7889 | 599 |
| Lautropia | K | 66.10 | 1.17 | 35 | 2 | 2210316 | 4822 | 547 |
| Ferrovaceae | N | 94.72 | 0.61 | 0 | 1 | 2846278 | 17342 | 338 |
| Neisseriaceae | N | 76.45 | 0.00 | 0 | 2 | 1064305 | 82838 | 45 |
| Kingella | K | 83.00 | 0.74 | 14 | 1 | 1552009 | 7513 | 305 |
| Kingella | N | 72.30 | 0.61 | 25 | 1 | 1669529 | 4626 | 436 |
| Neisseria | K | 89.25 | 1.87 | 30 | 2 | 2182285 | 11132 | 381 |
| Neisseria | N | 56.29 | 2.18 | 59 | 2 | 1357089 | 2358 | 632 |
| Neisseriaceae | K | 90.42 | 0.43 | 0 | 1 | 1302208 | 65201 | 44 |
| Pantoea | K | 60.85 | 2.59 | 0 | 1 | 3400613 | 7655 | 706 |
| Moraxella | K | 81.87 | 0.19 | 0 | 3 | 1517887 | 4484 | 437 |
| Moraxella | N | 75.23 | 0.41 | 0 | 2 | 1776873 | 6895 | 376 |
| Pseudomonas | N | 57.84 | 0.31 | 0 | 1 | 4264403 | 3273 | 1635 |
| Luteimonas | N | 78.91 | 1.92 | 0 | 1 | 2364043 | 8742 | 425 |
| Stenotrophomonas | K | 78.81 | 0.44 | 7 | 2 | 3767673 | 44793 | 653 |

^&^ Bacterial taxonomy at genus or the highest level of classification. ^#^K-known; N-novel MAGs.

**Table S5:** Summary of fungal metagenome-assembled genomes (MAGs) reconstructed from RAC/Face samples.

| **ID (blastn)** | **Drep clustering ID^&^** | **Class^#^** | **Genome Size** | **Completeness** |
| --- | --- | --- | --- | --- |
| Sordariomycetes | Sordariomycetes | N | 36508974 | 93.5 |
| Malasseziaceae | Malassezia globosa | K | 8801134 | 87.5 |
| Malassezia | Malassezia arunalokei | K | 7308028 | 86.3 |
| Malasseziaceae | Malassezia restricta | K | 5155029 | 66.5 |
| Malasseziaceae | Malasseziaceae | N | 6727485 | 83.1 |
| Malassezia | Malasseziaceae | N | 6446789 | 73.1 |

^&^ closest neighbor (refseq) within 97% ANI (fastANI). ^#^K-known; N-novel MAGs.

**Table S6:** Nested PERMANOVA results for Yanomami cutaneous microbiota (n=94) highlight factors that contribute to variation in bacterial and fungal community composition.

|  | **Body Site** | **Age Group** | **Expedition** | **Subject**  **(nested within Age group)** |
| --- | --- | --- | --- | --- |
| Model factor^*^ | Fixed | Fixed | Fixed | Random |
| **Bacteria^#^** | | | | |
| p(MC) | 0.0001 | 0.004 | 0.0004 | (no test) |
| Component of variation (sqrt) | 20.4% | 9.4% | 11.4% |  |
| *Interaction with* |  | **Body Site** | **Body Site** |  |
| p(MC) |  | 0.0007 | 0.0001 |  |
| Component of variation (sqrt) |  | 8.6% | 14.4% |  |
| **Fungi^#^** | | | | |
| p(MC) | 0.0001 | 0.0236 | 0.0021 | (no test) |
| Component of variation (sqrt) | 20.9% | 10.1% | 12.5% |  |
| *Interaction with* |  | **Body Site** | **Body Site** |  |
| p(MC) |  | 0.0001 | 0.0001 |  |
| Component of variation (sqrt) |  | 12.1% | 12.1% |  |

^#^Analysis performed on square root-transformed Bray-Curtis dissimilarity indices. ^*^PERMANOVA model: BC dissimilarity ~ Body Site + Age group + Expedition + Subject

p(MC) = Monte Carlo-simulated p-value.

**Table S7** – Taxa with ≥10% relative abundance in negative control sequencing reactions.

| **Kingdom** | **Strain** | **Kingdom** |  |
| --- | --- | --- | --- |
| **Bacteria/**  **Archaea** | *Bacillus sp. EGD-AK10* | **Fungi** | *Malassezia sp.* |
|  | *Cellulosimicrobium aquatile* |  | *Brettanomyces bruxellensis_u_t* |
|  | *Cellulosimicrobium cellulans LMG 16121* | **Phage** | *Lactococcus phage bIL310_u_t* |
|  | *Cellulosimicrobium sp. TH-20* |  | *Propionibacterium phage ATCC29399B_T* |
|  | *Cutibacterium acnes* |  | *Staphylococcus phage StB20-like_u_t* |
|  | *Propionibacterium sp. KPL1854* |  | *Stenotrophomonas phage phiSHP2_u_t* |
|  | *Stenotrophomonas maltophilia M30* |  | *Stenotrophomonas phage Smp131_u_t* |
|  | *Stenotrophomonas sp. PAMC25021* | **Protist** | *Pseudoperonospora cubensis* |
| **Fungi** | *Malassezia globosa CBS 7966* | **Virus** | *Merkel cell polyomavirus* |
|  | *Malassezia restricta* |  |  |

**Supplemental Figures:**

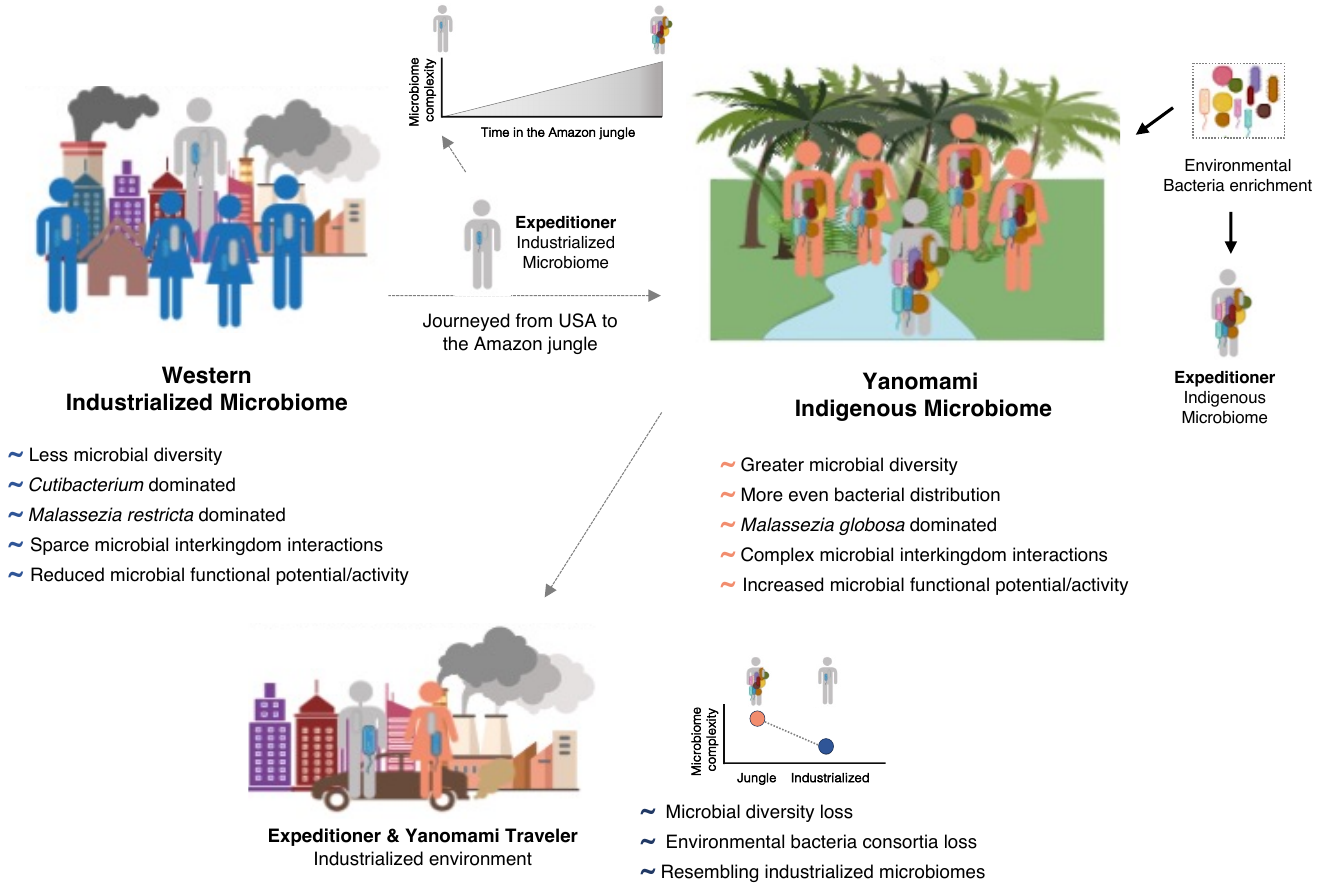

**Figure S1:** A schematic of study design highlighting major findings.

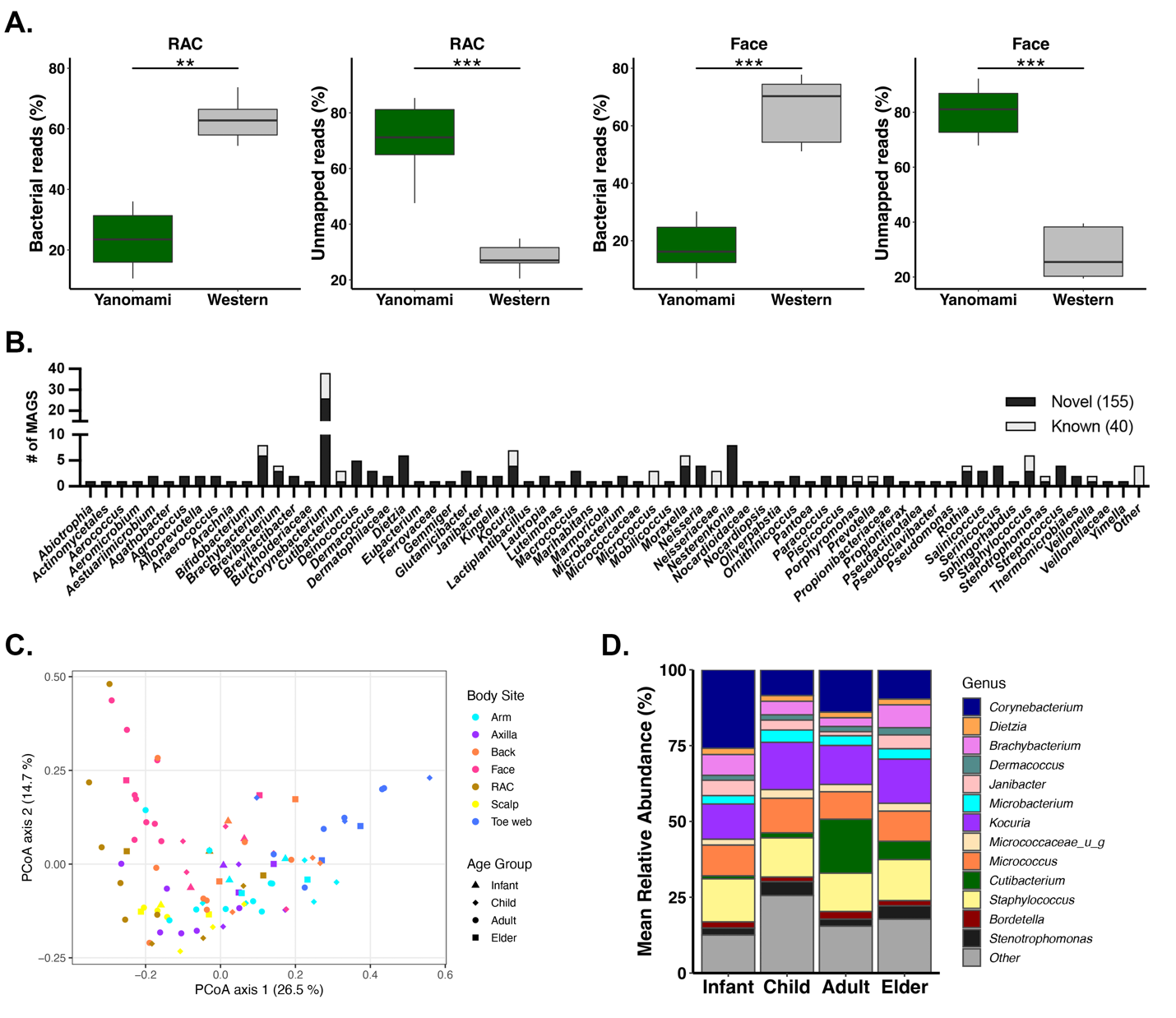

**Figure S2:** **A.** Percent of rarefied reads (13M) that mapped to bacterial genomes or were unmapped in Yanomami RAC (n=12) or face (n=18) samples compared to western expeditioner (n=5 each body site). ANOVA of LME p-value: ***<0.001, **<0.01. **B.** Classification of known and novel bacterial MAGs reconstructed from Yanomami RAC/Face samples (n=30) classified using The Skin Microbial Genome Collection (SMGC). **C.** Principal Coordinates Analysis of Bray-Curtis dissimilarity indices on cutaneous mycobiota of the Yanomami (n=94). **D.** Relative abundance of bacterial genera (≥2%) in sebaceous samples (RAC/Face/Back/Scalp; n=53) in individuals grouped by age.

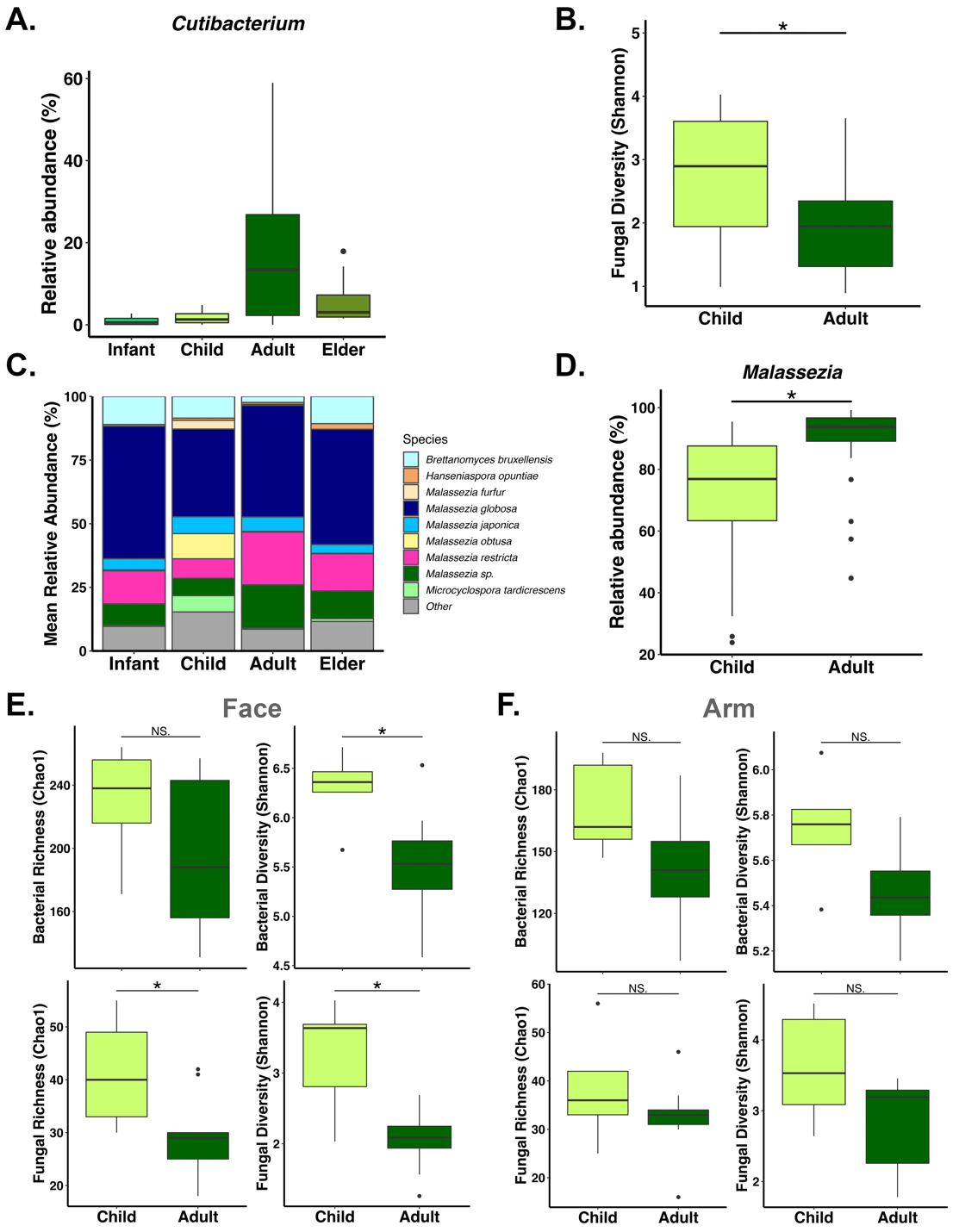

**Figure S3:** **A.** Relative abundance of *Cutibacterium* in Yanomami individuals’ combined sebaceous samples (RAC/Face/Back/Scalp; n=53) by age group (ANOVA of LME p=0.12). **B.** Fungal Diversity (Shannon Index) of combined sebaceous samples (RAC/Face/Back/Scalp; n=40) in Yanomami individuals grouped by age. (ANOVA of LME p-value: *<0.05). **C.** Relative abundance of fungal species (≥2%) in combined sebaceous samples (n=53) by age group. **D.** Relative abundance of *Malassezia* in combined sebaceous samples (n=40) by age group. (ANOVA of LME p-value: *<0.05). **E.** Bacterial and Fungal richness and diversity on the face (a sebaceous skin site) of Yanomami children (n=5) and adults (n=9; Wilcoxon test p-value: *<0.05, NS >0.05). **F.** Bacterial and Fungal richness and diversity on the arm (a dry skin site) of Yanomami children (n=5) and adults (n=9; Wilcoxon test p-value: NS >0.05).

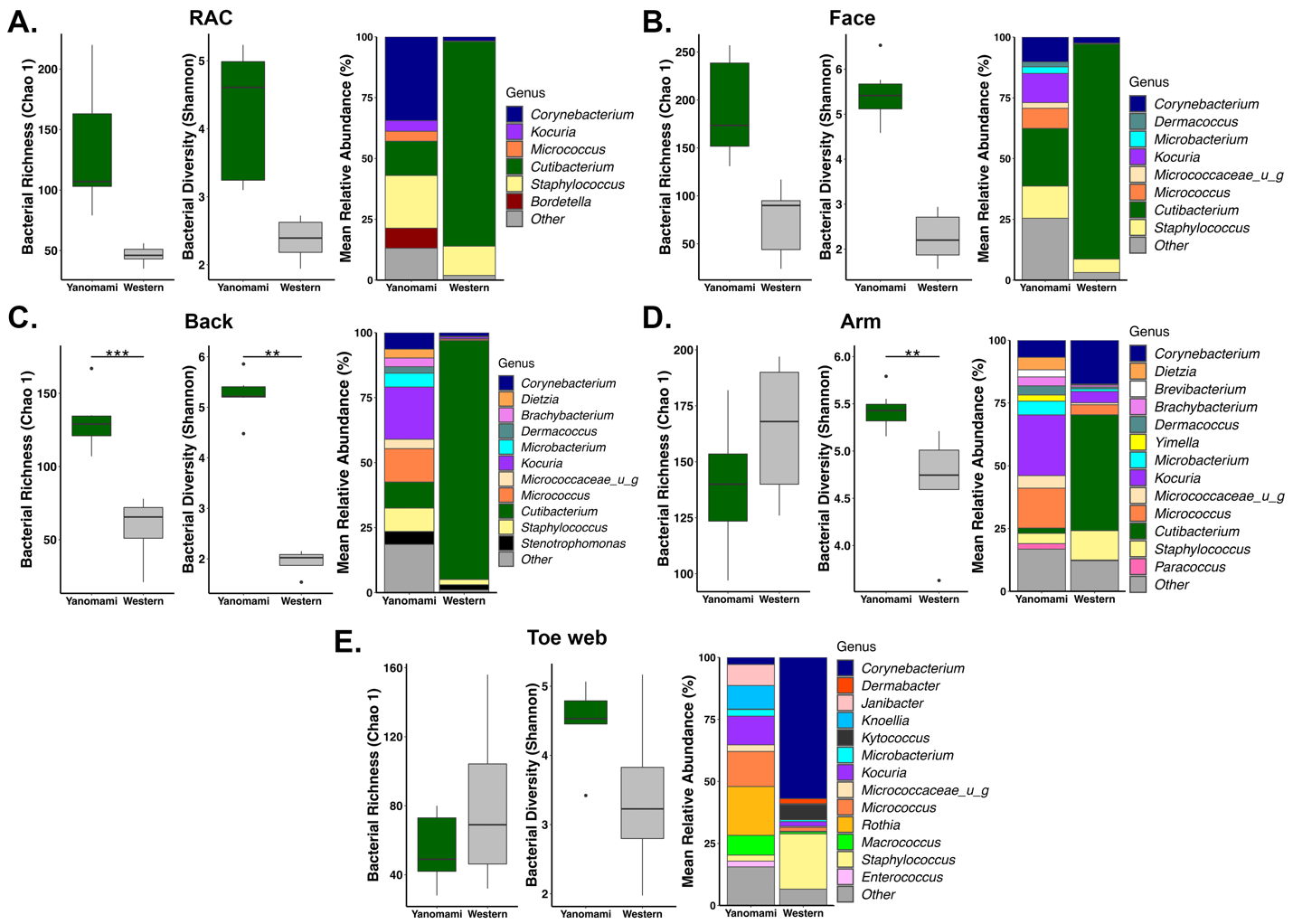

**Figure S4:** Bacterial alpha diversity (Chao1 index – richness; Shannon index – diversity) and mean relative abundance of genera (≥2%) at distinct body sites of the Yanomami adults compared to western expeditioner collected in the USA or in transit (i.e., not in Amazon): **A.** RAC (n=5; n=5), **B.** Face (n=8, n=5), **C.** Back (n=7, n=4), **D.** Arm (n=8, n=5), **E.** Toe web (n=5, n=4), respectively (ANOVA of LME: *** p<0.001, ** p<0.01).

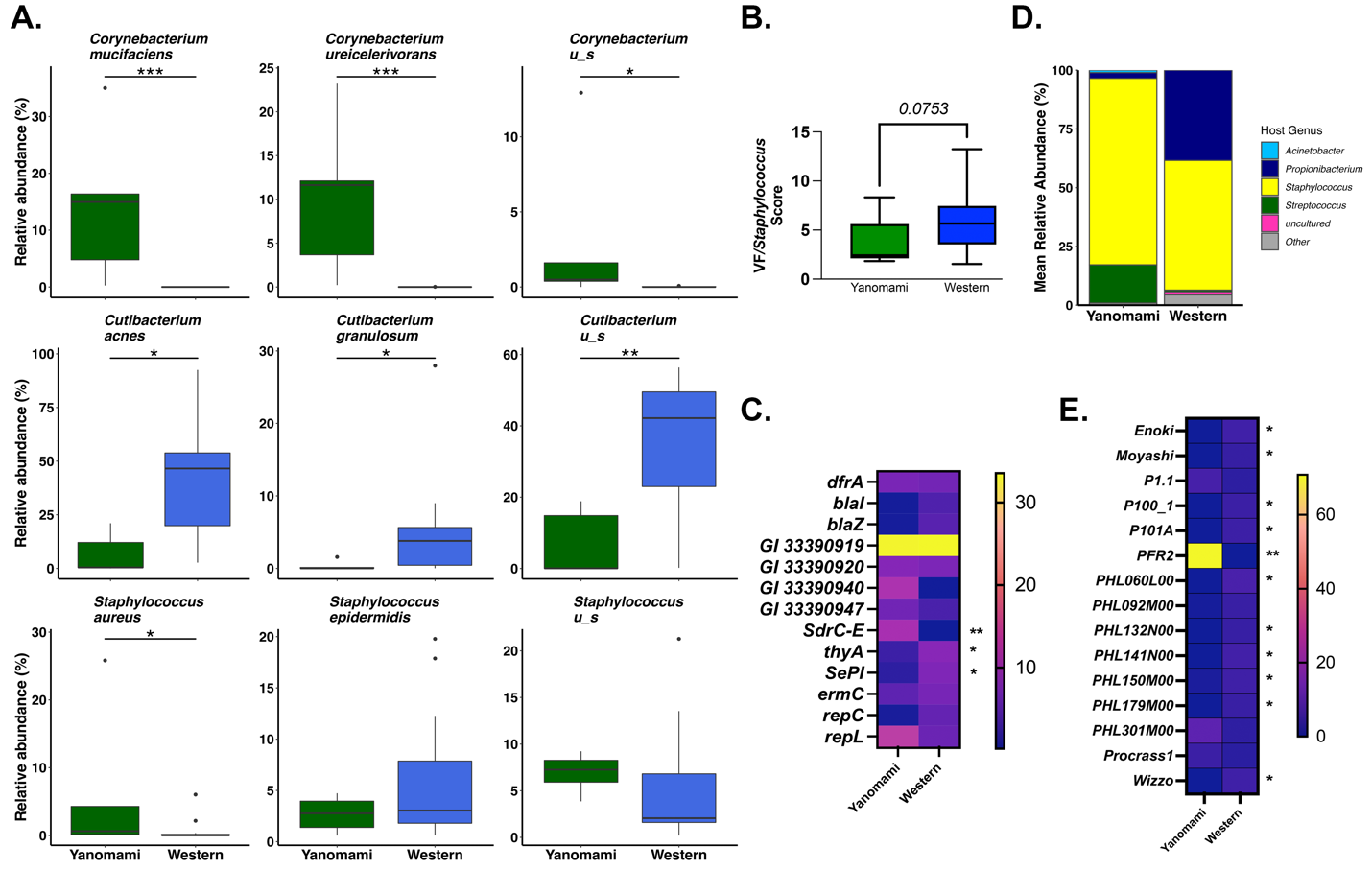

**Figure S5:** **A.** Relative abundance of most abundant bacterial species that are similarly frequently detected in RAC samples from Yanomami adults (n=5) and HMP (n=16) westerners. (Wilcoxon test with Benjamini-Hochberg correction: *** q <0.001, ** q<0.01, * q<0.05). **B.** Ratio of combined *Staphylococcus* virulence factors to *Staphylococcus* abundance score (Wilcoxon test) in RAC samples from Yanomami adults (n=5) and HMP (n=16) westerners. **C.** Mean abundance score of *Staphylococcus* virulence factors (at ≥2%; Wilcoxon test with Benjamini-Hochberg correction: **** q<0.001, *** q<0.01, ** q<0.05, * q<0.1) in RAC samples from Yanomami adults (n=5) and HMP (n=16) westerners. **D.** Mean relative abundance of bacteriophage (≥1%) in ≥25% samples in ≥1 population, grouped by bacterial host in RAC samples from Yanomami adults (n=5) and HMP (n=16) westerners. **E.** Mean relative abundance of *Propionibacterium*-associated phage species (≥2%, based on total *Propionibacterium*-associated phage abundance score) in RAC samples from Yanomami adults (n=5) and HMP (n=16) westerners. (Wilcoxon test with Benjamini-Hochberg correction, ** q<0.01, * q<0.05).

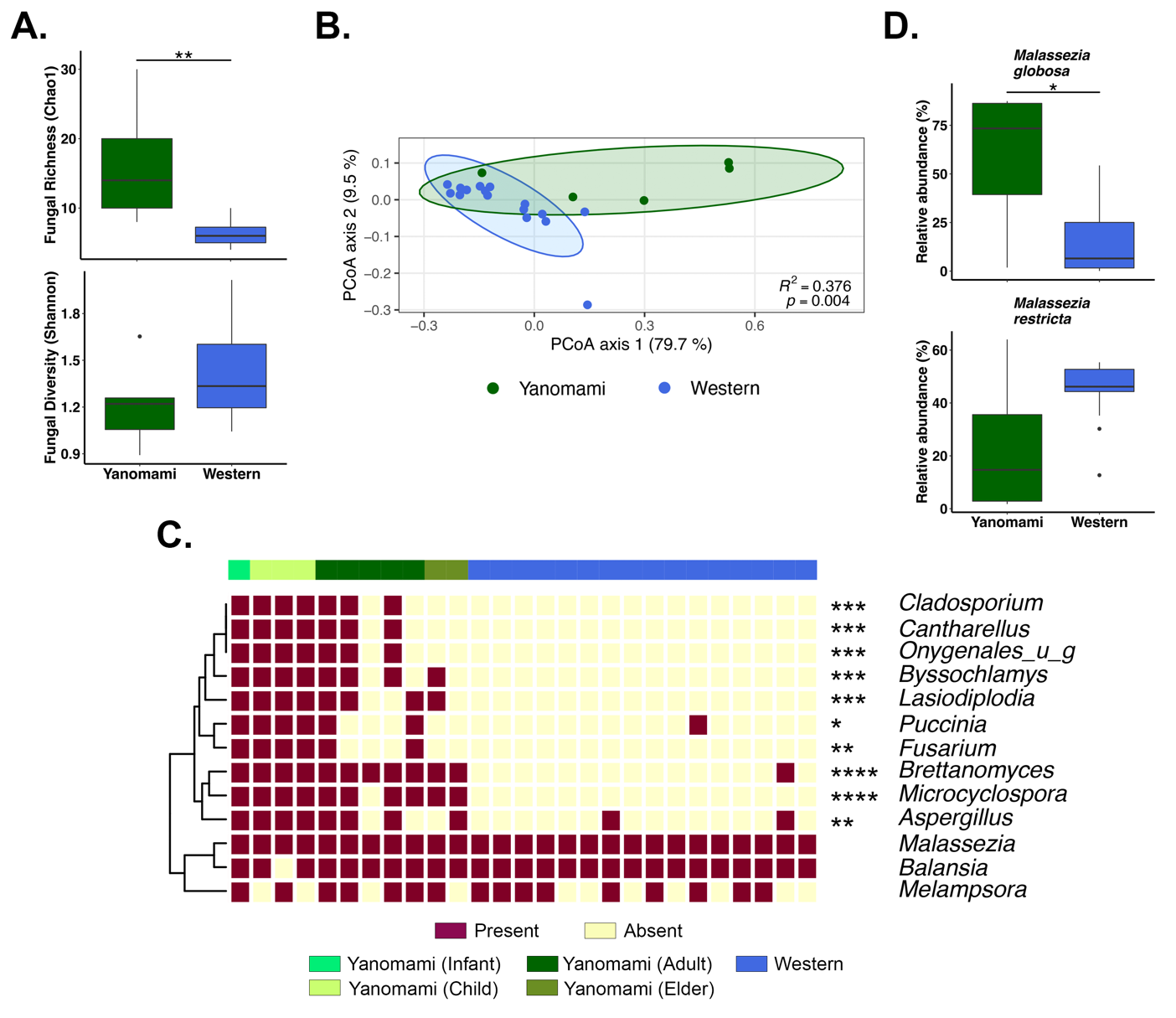

**Figure S6:** **A.** Fungal Richness (Chao1 index) and Diversity (Shannon Index) in RAC samples of Yanomami adults (n=5) and HMP (n=16) westerners (Wilcoxon test p-value: ** p<0.01, not shown p>0.05). **B.** Fungal community composition (PERMANOVA of Bray-Curtis dissimilarity) in RAC samples of Yanomami adults (n=5) and HMP (n=16) westerners. **C.** Distribution of fungi at >50% prevalence in either Yanomami RAC samples (n=11; colored by age group) or HMP (n=16) westerners (Fisher’s exact test, FDR-adjusted: **** q<0.0001, *** q<0.001, ** q<0.01, * q<0.05). **D.** Relative abundance of most abundant yeast species that are similarly frequently detected in RAC samples from Yanomami adults (n=5) and HMP (n=16) westerners (Wilcoxon test with Benjamini-Hochberg correction: * q<0.05).

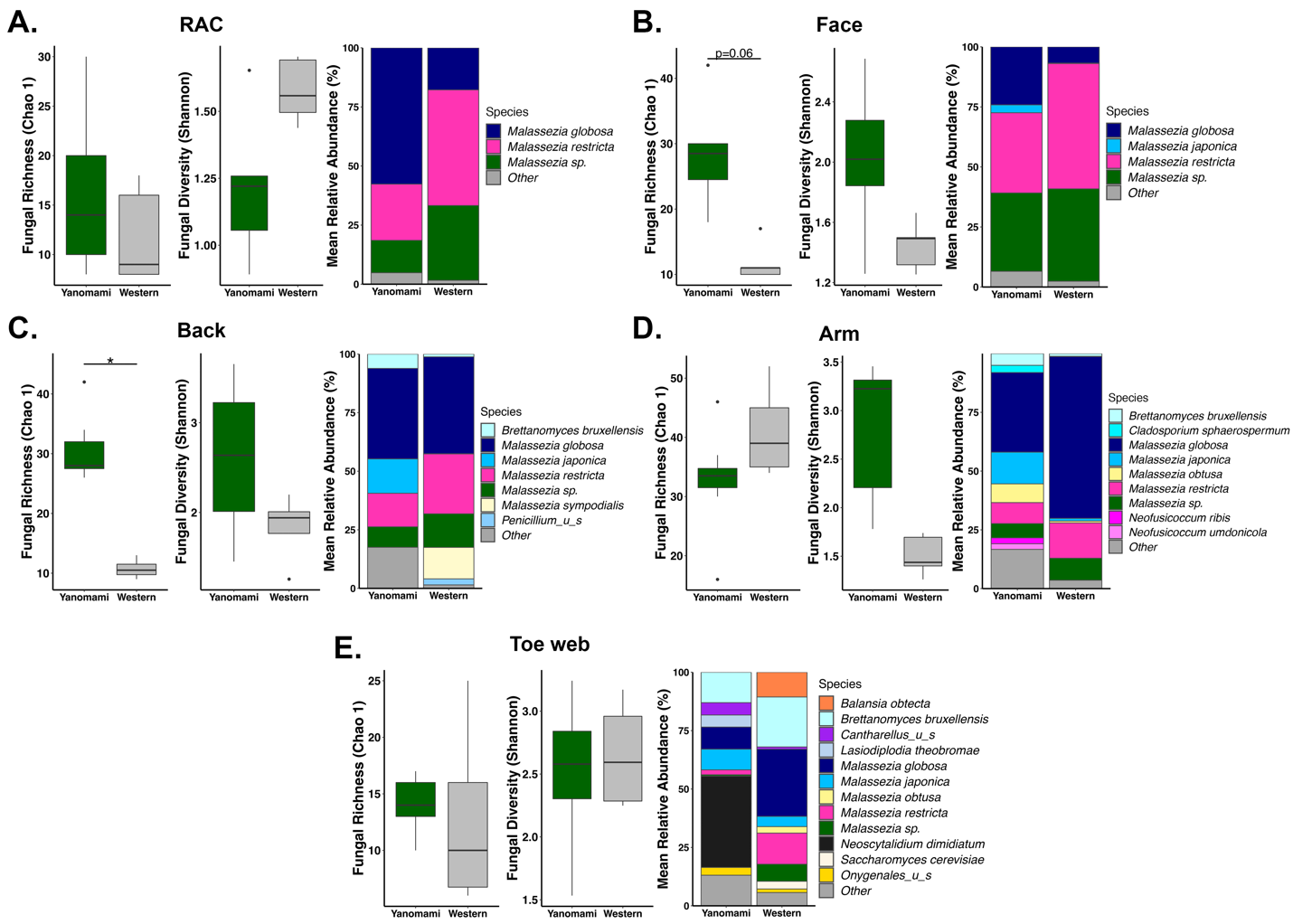

**Figure S7:** Fungal alpha diversity (Chao1 index – richness; Shannon index – diversity) and mean relative abundance of species (≥2%) at distinct body sites of the Yanomami adults compared to western expeditioner collected in the USA or in transit (i.e., not in Amazon): **A.** RAC (n=5; n=5), **B.** Face (n=8, n=5), **C.** Back (n=7, n=4), **D.** Arm (n=8, n=5), **E.** Toe web (n=5, n=4), respectively (ANOVA of LME: * p<0.05).

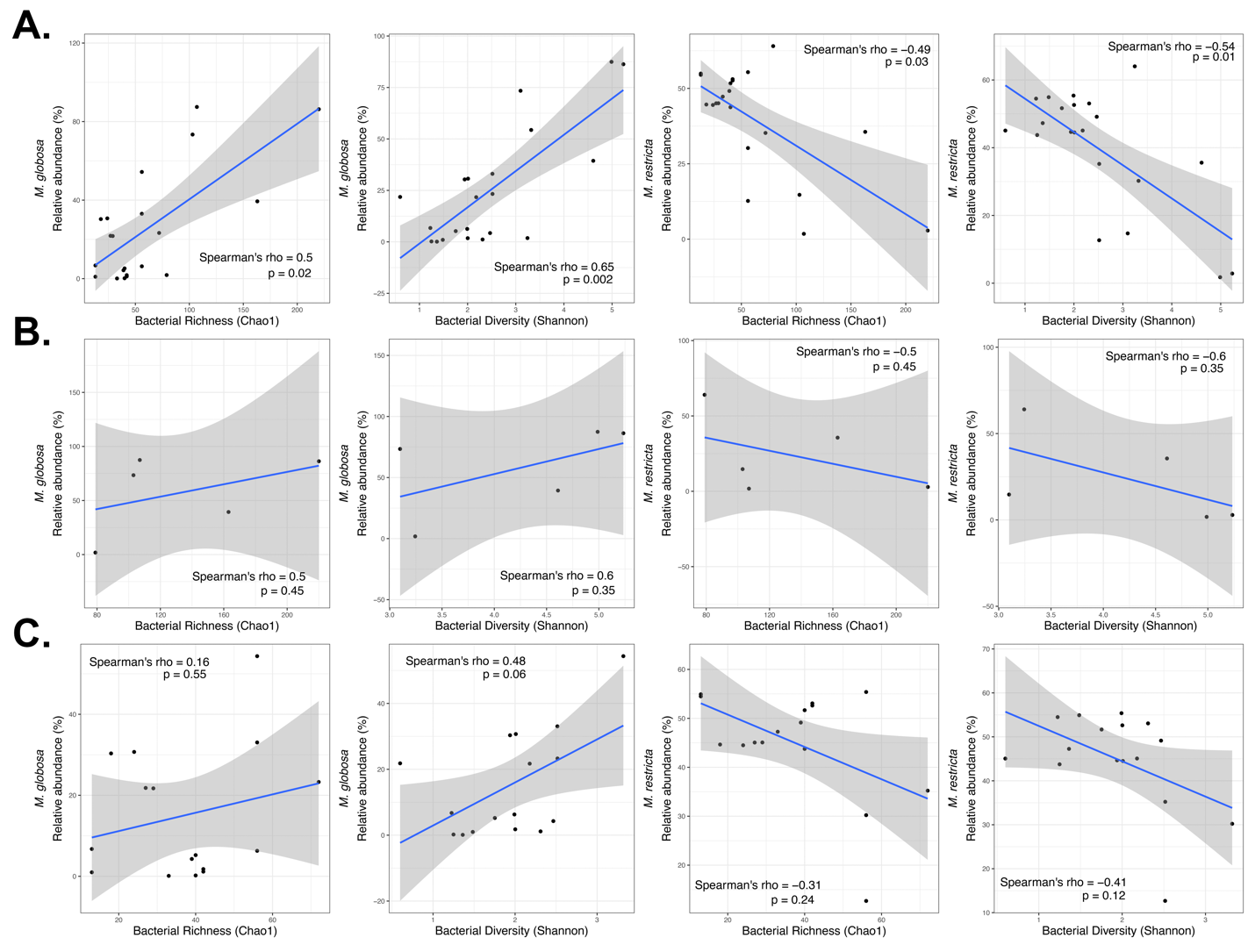

**Figure S8:** Linear regression plots showing correlation between relative abundance of *Malassezia globosa* or *M. restricta* versus bacterial richness or diversity in RAC samples of **A.** adult Yanomami and HMP western individuals (n= 21), **B.** adult Yanomami (n=5), **C.** HMP westerners (n=16).

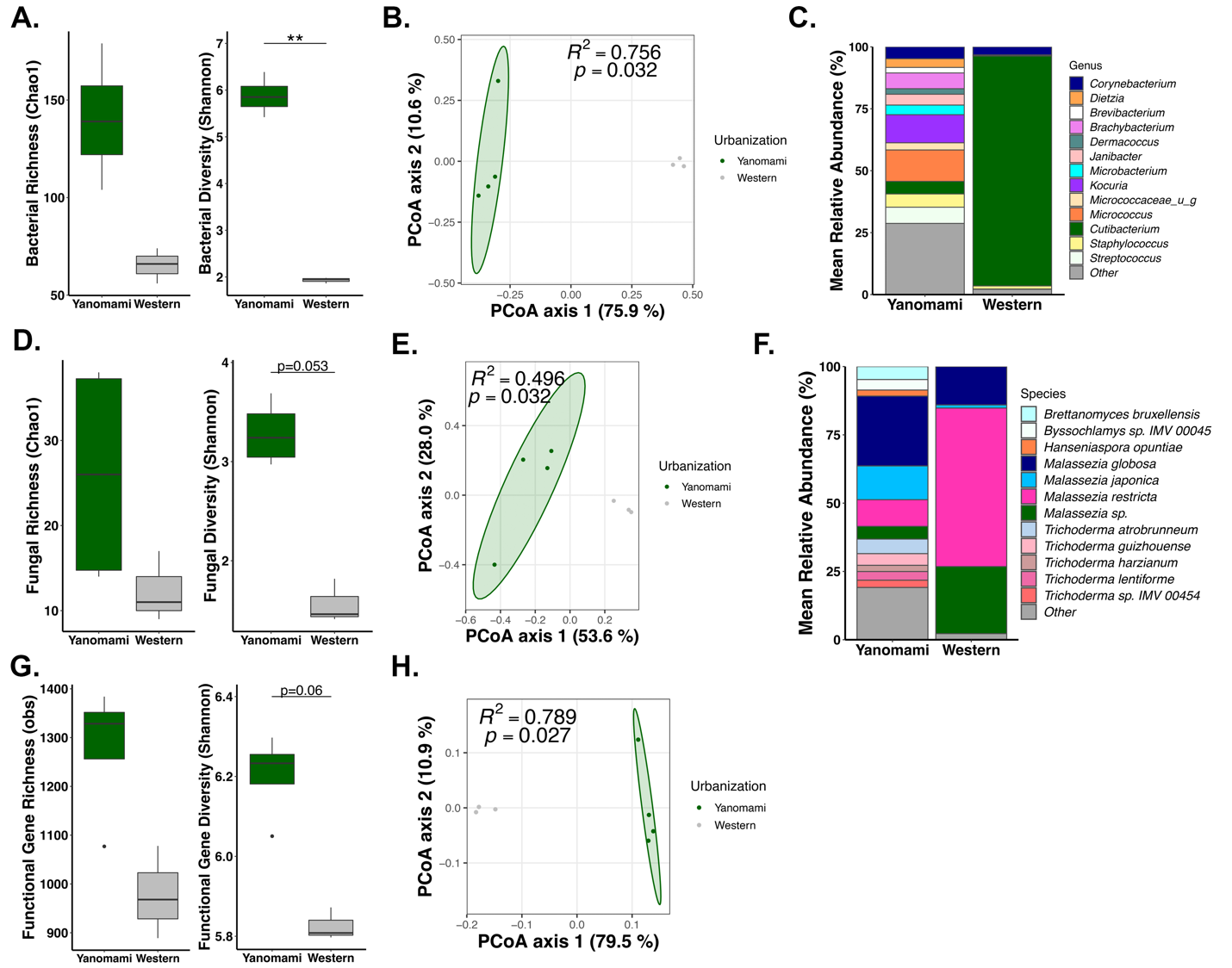

**Figure S9:** **A.** Bacterial Richness (Chao1 Index) and Diversity (Shannon Index) of active microbes (RNA) in Yanomami adult (n=4) and western expeditioner (n=3, non-Amazon; ANOVA of LME: **<0.01) chest samples. **B.** Active bacterial community composition (RNA, PERMANOVA of Bray-Curtis dissimilarity on species data) in Yanomami adult (n=4) and western expeditioner (n=3, non-Amazon) chest samples. **C.** Mean relative abundance of active bacterial genera (≥2%) in Yanomami adult (n=4) and western expeditioner (n=3, non-Amazon) chest samples. **D.** Fungal Richness (Chao1 Index) and Diversity (Shannon Index) of active microbes (RNA) in Yanomami adult (n=4) and western expeditioner (n=3, non-Amazon; ANOVA of LME) chest samples. **E.** Active fungal community composition (RNA, PERMANOVA of Bray-Curtis dissimilarity on species data) in Yanomami adult (n=4) and western expeditioner (n=3, non-Amazon) chest samples. **F.** Mean relative abundance of active fungal species (≥2%) in Yanomami adult (n=4) and western expeditioner (n=3, non-Amazon) chest samples. **G.** Richness (obs) and Diversity (Shannon index) of active microbiota functional pathways (pathways assigned by MetaCyc) of the Yanomami (chest, n=4) and western expeditioner (chest, n=3) skin microbiome. Data were normalized to counts per million (CPM; ANOVA of LME: ***<0.001). **H.** Functional pathways composition (RNA, PERMANOVA of Bray-Curtis dissimilarity; pathways assigned by MetaCyc) in Yanomami adult (n=4) and western expeditioner (n=3, non-Amazon) chest samples.

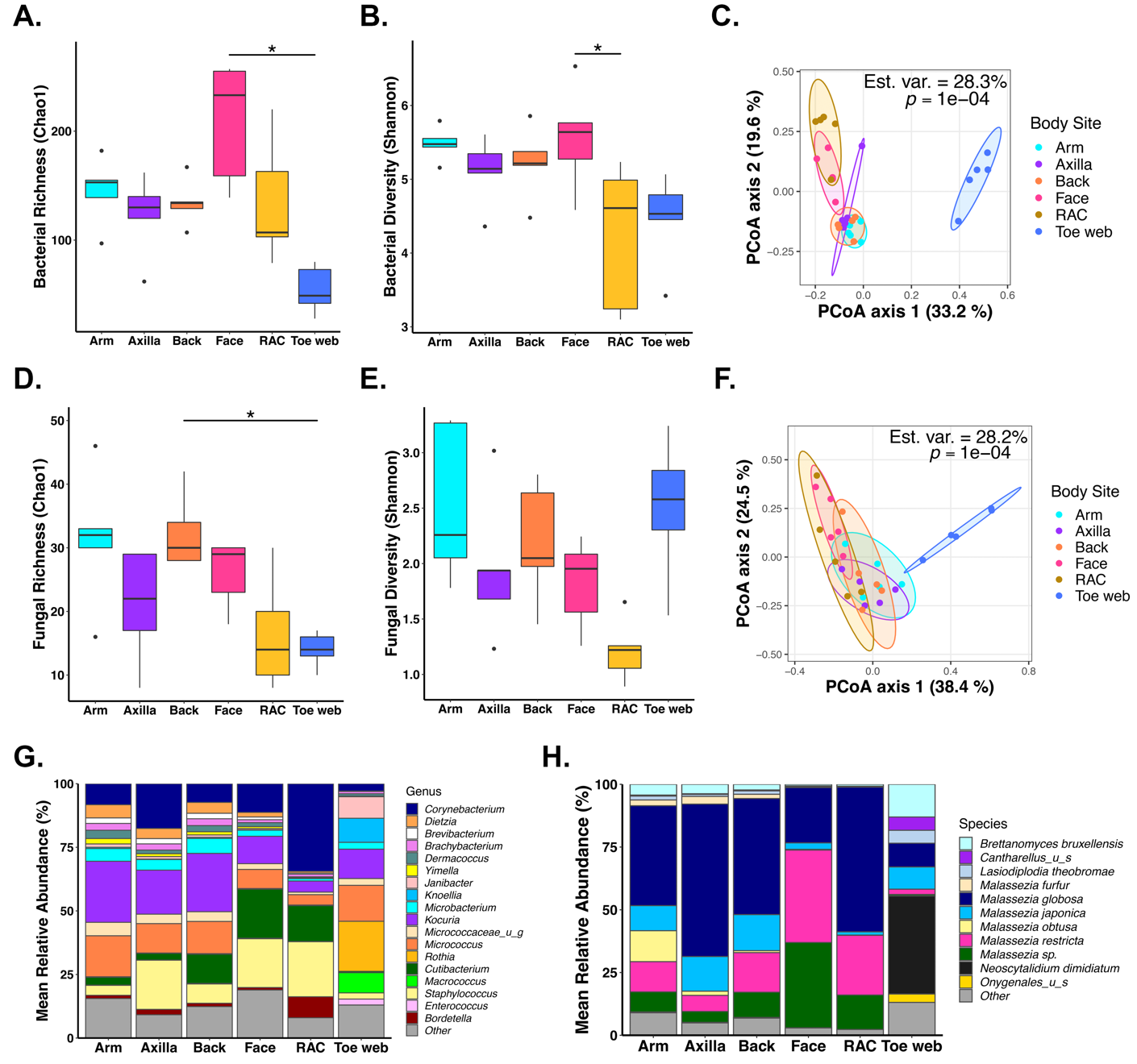

**Figure S10:** **A.** Bacterial species richness (Chao1 index; Friedman [group-wise] p<0.01, Dunn test [pair-wise, BH-corrected p<0.025]), **B.** diversity (Shannon index; Friedman [group-wise] p<0.01, Dunn test [pair-wise, BH-corrected p<0.025]), and **C.** community composition (PERMANOVA of Bray-Curtis dissimilarity) by body site of Yanomami adults (n=5). **D.** Fungal species richness (Chao 1 index, Friedman [group-wise] p=0.04, Dunn test [pair-wise, BH-corrected p<0.025]), **E.** diversity (Shannon index, Friedman [group-wise] p=0.06), and **F.** community composition (PERMANOVA of Bray-Curtis dissimilarity) by body site of Yanomami adults (n=5). **G.** Relative abundance of bacterial genera (≥2%) and **H.** fungal species (≥2%) by body site of Yanomami adults (n=5).

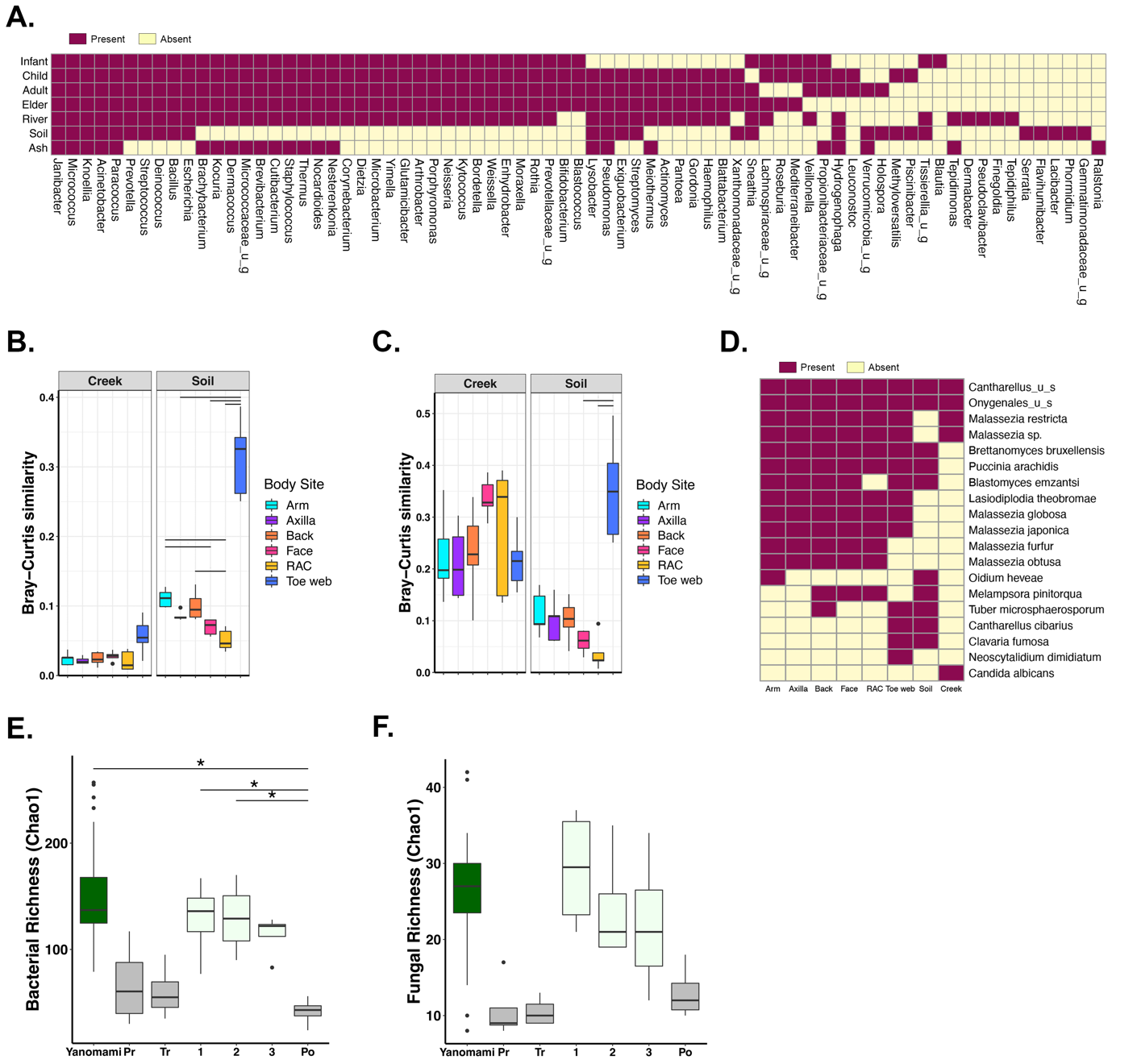

**Figure S11:** **A.** Distribution of active bacterial genera with ≥0.5% relative abundance in Yanomami chest samples, by age group (Infant [n=2], Child [n=4], Adult [n=4], Elder [n=2]), and/or environmental samples. **B.** Community compositional similarity (Bray-Curtis) of adult Yanomami (n=5 adults) skin microbiota at distinct body sites to their environment (Friedman [group-wise]: Creek p>0.05, Soil p<0.001, Dunn test [pair-wise, BH-corrected p<0.025]) or **C.** skin mycobiota to same (Friedman [group-wise]: Creek p>0.05, Soil p<0.01, Dunn test [pair-wise, BH-corrected p<0.025]). **D.** Distribution of fungal species with ≥2% relative abundance in Yanomami adults (n=5) across various body sites or at any percentage in environmental samples. **E.** Bacterial species richness (ANOVA of LME p=0.044, Tukey multiple comparisons: *** p<0.001; ** p<0.01; * p<0.05) and **F.** Fungal species richness (ANOVA of LME p=0.180) of adult Yanomami (n=9) combined sebaceous (back, face, RAC, scalp) sites (n=26 samples) compared with the expeditioner sampled longitudinally (same body sites) during the 2018 expedition (n=23 samples; Pr - pre-expedition, Tr - in-transit, 1 – first sample in Amazon, 2 – second Amazon sample, 3 – third Amazon sample, Po – post-expedition).

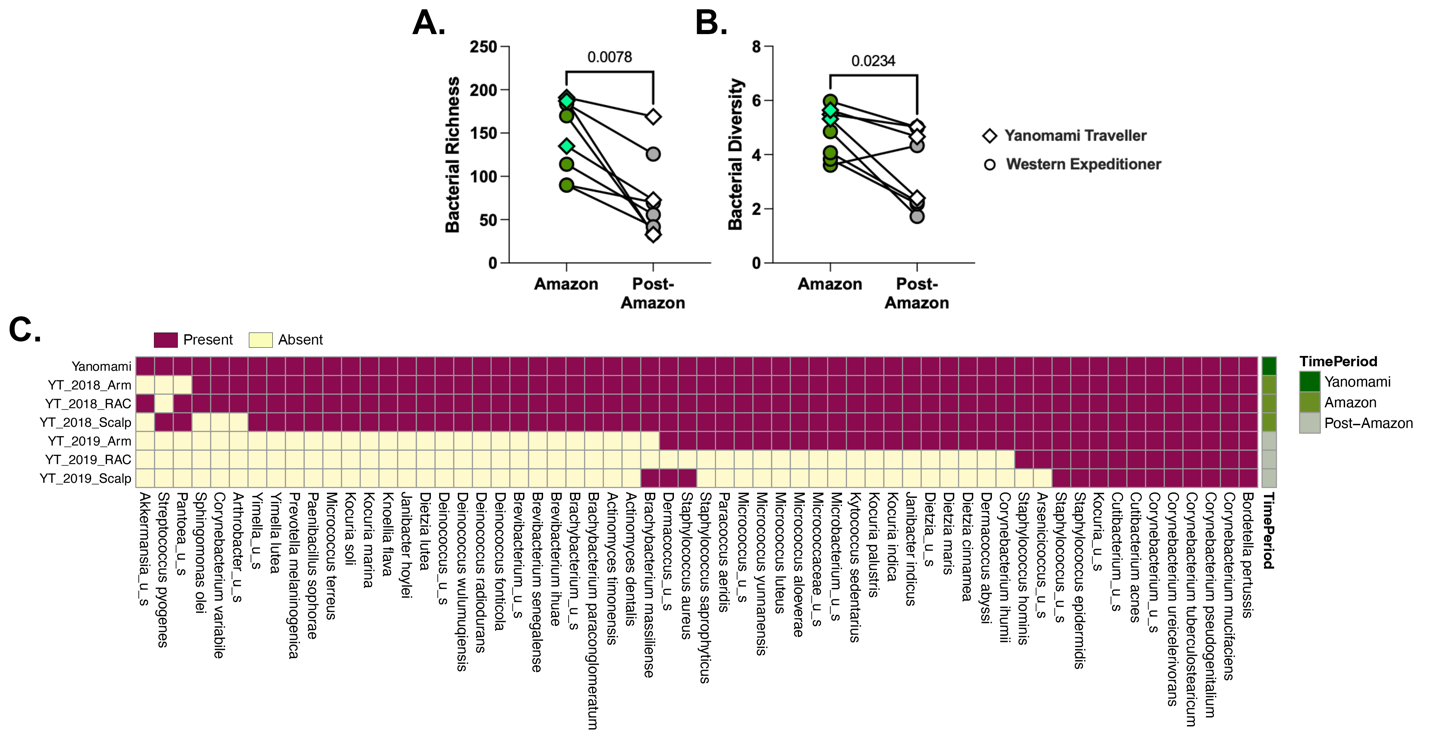

**Figure S12:** richness (Chao1 index) and **B.** diversity (Shannon index) of paired samples from various body sites of travelers (western expeditioner - samples included arm, axilla, face, RAC, and scalp or Yanomami traveler [YT] samples – arm, RAC, scalp) while in the Amazon or following westernization (paired Wilcoxon test). **C.** Distribution (presence/absence) of bacterial species lost from YT following westernization, selected to meet the following criteria: prevalent on the Yanomami (present in ≥25% samples at ≥0.5% relative abundance [average of n=80, toe web samples excluded]), having ≥0.25% relative abundance on the Yanomami and absent in YT samples following westernization, or having ≥0.25% relative abundance on YT (2018 - in the Amazon) and absent on YT [2019 - following westernization].
