## Supplemental Methods for "Yanomami skin microbiome complexity challenges prevailing concepts of healthy skin"

**Extended Methods:**

**Sample Collection**

Cutaneous microbiome samples were collected using sterile, viscose rayon swabs (Isohelix, Kent, UK) and then preserved in DNA/RNA Shield (Zymo Research, Irvine, CA). Swabs were moistened with sterile PBS prior to sample collection. The following body sites were sampled for 30 seconds by a gloved operator: arm (antecubital fossa and volar forearm, combined), axilla, back (upper area), chest, face (forehead, cheeks, nose, alar crease, and chin, combined), RAC (retroauricular crease), scalp, toe web. Preserved samples were stored at -80°C within 7 weeks of collection. Environmental samples were collected using swabs (Expedition1 – hearth ash, soil, water; Expedition2 - soil) or Sterivex filters (Expedition2, water). For the latter, 30 mL water were collected into a sterile syringe and then filtered through a Sterivex filter. Subsequently, DNA/RNA Shield was added to the Sterivex filter cartridge and end caps affixed to preserve the sample.

**Nucleic acid extraction and metagenomic sequencing**

DNA was extracted from back swabs using ZymoBIOMICs 96 MagBead DNA Kit (Zymo Research, Irvine, CA, USA). MetaPolyzyme (Sigma-Aldrich, St. Louis, MO, USA) pre-treatment [6 uL enzyme added to samples, incubated at 35°C for 16 hr] followed by ZymoBIOMICS Microprep kit was used to extract DNA for the remaining body site samples (swabs). Qiagen PowerSoil Pro kit (Qiagen, Germantown, MD, USA) was used to extract DNA from soil and ¼ filters (2019 water samples). RNA was extracted from chest and environmental samples (swabs) using the Qiagen RNeasy Plus kit (Qiagen, Germantown, MD, USA). DNA was sequenced using the Nextera XT library prep kit (Illumina, San Diego, CA, USA). RNA was sequenced using the SMARTer Stranded RNA Total RNA Sample Prep Kit - Pico Input (Takara Bio USA, San Jose, CA, USA). All sequencing was performed by CosmosID using an Illumina HiSeq 4000 instrument and paired-end, 2 x 150 cycle reagent kit (Illumina, San Diego, CA, USA).

**Data analysis**

Sequence reads were demultiplexed through BaseSpace (Illumina, San Diego, CA, USA), and quality parameters were evaluated using FastQC and MultiQC. No quality-trimming was performed before taxonomic analysis. Human reads were removed using CosmosID’s kmer-based host-read removal pipeline (using genome assembly GRCh38.p6). Percentages of reads mapped to each taxonomic kingdom were calculated prior to and following normalization to 13M reads per sample. Samples were normalized using reformat.sh of the BBTools suite^1^. The output fasta files were then analyzed through CosmosID-HUB portal.

*Taxonomic Classification*

The CosmosID-HUB uses a high-performance data-mining k-mer algorithm that rapidly disambiguates millions of short sequence reads into the discrete genomes comprising the particular sequences. The pipeline has two separable comparators: the first consists of a pre-computation phase for reference databases and the second is a per-sample computation. The input to the pre-computation phase are databases of reference genomes, virulence markers, and antimicrobial resistance markers that are continuously curated by CosmosID scientists. The output of the pre-computational phase is a phylogeny tree of microbes, together with sets of variable length k-mer fingerprints (biomarkers) uniquely associated with distinct branches and leaves of the tree. The reference database comprises publicly available genomes or gene sequences through NCBI- RefSeq/WGS/SRA/nr, PATRIC, M5NR, IMG, ENA, DDBJ, CARD, ResFinder, ARDB, ARG-ANNOT, mvirdb, VFDB etc., as well as a subset of genomes sequenced by CosmosID and its collaborators. The second, per-sample computational phase searches the hundreds of millions of short sequence reads, or alternatively contigs from draft de novo assemblies, against the fingerprint sets. This query enables the sensitive and highly precise detection and taxonomic classification of microbial NGS reads. The resulting statistics are analyzed to return the fine-grain taxonomic and relative abundance estimates for the microbial NGS datasets. To exclude false positive identifications, the results are filtered using a threshold based on internal statistical scores that are determined by analyzing a large number of diverse metagenomes. The same approach is applied to enable the sensitive and accurate detection of genetic markers for virulence and for resistance to antibiotics. Reads were assigned to bacterial & archaeal, fungal, protist, or viral (excluding phage), or bacteriophage lineages for taxonomic comparisons.

Taxa representing ≥10% relative abundance in ≥1 negative extraction control(s) were identified as likely/possible extraction and/or sequencing contaminants (Table S6) but not removed because the relative abundance of some taxa was much greater in samples than in controls and because several taxa are plausible cutaneous or environmental microbes. Two bacterial sequences included in a DNA and the RNA positive extraction control samples (*Allobacillus halotolerans,* *Imtechella halotolerans;* ZymoBIOMICS Spike-in Control I, (D6320, Zymo Research, Irvine, CA, USA)) were removed from the dataset prior to analysis. The ZR ZymoBIOMICS Microbial Community Standard (D6300) was used as a positive control in some extraction sets. This standard contains organisms plausibly in cutaneous or environmental samples, so they were not removed.

Previously published data from Human Microbiome Project (HMP) cutaneous RAC samples (n=16, rarefied to 13M reads, with the following NCBI Short Read Archive sample identifiers SRS013261, SRS020263, SRS045606, SRS047134, SRS050557, SRS058221, SRS058646, SRS075093, SRS075356, SRS077864, SRS078257, SRS078434, SRS078741, SRS100036, SRS101448, SRS102817, which were right RAC samples with >1.5 GB data, except when only a left sample had ≥13M de-hosted reads) were used for comparative analyses. Although human reads had already been removed from the available data files, the data were processed through the same data analysis pipeline, including human read removal, to assure the most direct comparisons.

*Functional Classification*

Initial QC, adapter trimming and preprocessing of metagenomic sequencing reads are performed using BBduk^1^. Human reads were removed (de-hosted) using ‘bmtagger’ (ftp://ftp.*ncbi*.nlm.nih.gov/pub/agarwala/*bmtagger*/) with hg38 as the human reference genome. The quality-controlled reads were analyzed without down-sampling (RNA-based analyses) or were down-sampled to 13 million reads (DNA-based analyses) using BBTools suite^1^. Both sets of quality-controlled reads were subjected to a translated search against a comprehensive and non-redundant protein sequence database, UniRef 90. The UniRef90 database, provided by UniProt^2^, represents a clustering of all non-redundant protein sequences in UniProt, such that each sequence in a cluster aligns with 90% identity and 80% coverage of the longest sequence in the cluster. The mapping of metagenomic reads to gene sequences are weighted by mapping quality, coverage, and gene sequence length to estimate community-wide weighted gene family abundances^3^. Gene families were then annotated to MetaCyc^4^ reactions (Metabolic Enzymes) before reconstructing and quantifying MetaCyc reactions to MetaCyc pathways in the community, as previously described^3^. Lastly, to facilitate comparisons across multiple samples with different sequencing depths at the MetaCyc pathways and MetaCyc reactions classifications, the abundance values are processed using Total-Sum Scaling (TSS) normalization to produce "Copies per million" (analogous to TPMs in RNA-Seq) units. The resulting metabolic reactions before normalizing are regrouped using a custom mapping table using the “regroup_table” function on HUMAnN3 to the MetaCyc “pathways” classification and normalized after. The resulting functional pathways table is “regrouped” into a functional table consisting of MetaCyc secondary pathway classes for analysis with the Linear discriminant analysis Effect Size (LEfSe)^5^ software package to determine significant differential enrichment of functions between Western and Yanomami samples.

*Metagenome assembly and taxonomy assignment*

Metagenome assembled genomes (MAGs) were assembled with ‘MetaWRAP’^6^ from face and RAC samples. Adapters were trimmed using ‘Trim Galore’ (https://github.com/FelixKrueger/TrimGalore) and ‘cutadapt’^7^. Human reads were removed (de-hosted) using ‘bmtagger’ (ftp://ftp.*ncbi*.nlm.nih.gov/pub/agarwala/*bmtagger*/) with hg38 as the human reference genome. Reads were assembled primarily with metaSPAdes^8^, then unassembled reads from metaSPAdes were assembled with MEGAHIT^9^. Assembled contigs were indexed and aligned to reads using SAMtools^10^ and BWA-MEM^11^. Bacterial contigs were then binned using MetaBAT2^12^, MaxBin 2^13^, and CONCOCT^14^. Bin quality was assessed, and the best bin chosen using the refine command from MetaWRAP. Refined bins were further improved using the reassemble command from MetaWRAP, which uses metaSPAdes to remap the de-hosted reads to each of the bins. To assist bin refinement, checkM^15^ was used to assess the bin quality among the binners and select improved bins. MAGs with ≥50% completion and ≤10% contamination were retained. Next, dRep^16^ was used to dereplicate MAGs and provide a representative set of MAGs using the “-comp 50 -con 5 -pa 0.9 -sa 0.95 -nc 0.30 -cm larger” parameters. Distances among genomes were evaluated with a primary clustering threshold of 90% using MASH^17^ and secondary clustering threshold of 95% using FastANI^18^. Representative genomes had ≥50% completion and ≤5% contamination, following MIMAG standards^19^. Representative MAGs (n=195) were classified with GTDB-tk (v2.10 07-RS207^20^), which is a comprehensive database of GenBank isolate genomes, single-cell assembled genomes (SAGs), and MAGs. Representative MAGs from the Yanomami metagenomes that were within 95% average nucleotide identity (ANI) to a genome from the GTDB are classified as “known”. Representative MAGs below 95% ANI or those that did not have an ANI calculated due to high sequence divergence were classified as “novel”. To compare representative MAGs from the Yanomami metagenomes against existing human skin MAG databases, the representative MAGs were also compared against the Skin Microbial Genome Collection^21^, using the same ANI criteria as used with GTDB.

Fungal contigs assembled from metaSPAdes were filtered from the metagenomic assembled contigs using EukRep^22^, indexed and aligned to reads using SAMtools and BWA-MEM, then binned using MetaBAT2. Bin quality was assessed using BUSCO^23^ with the fungi ortholog dataset. Fungal MAGs with ≥50% BUSCO completion were retained. Next, dRep was used to dereplicate fungal MAGs and provide a representative set of MAGs using the “-pa 0.9 -sa 0.95 -nc 0.30 -cm larger” parameters. Fungal MAGs were then classified using taxator-tk^24^ and Blast+^25^. Using the secondary clustering (fastANI), species that were within the threshold of 95% ANI of a NCBI assembly were classified as known and named per NCBI reference.

*Network Analysis*

Spearman’s correlation was calculated with ‘Hmisc’^26^ to analyze microbial abundance scores and detect microbial interactions using read-normalized data. In the CosmosID-HUB app, abundance score is a normalized, absolute abundance metric considering genome size and the number of reads. Taxa (bacterial genera and any Malassezia spp.) included in the analysis had ≥80% prevalence in adult Yanomami (n=5) and/or HMP (as western, n=16) samples. *Malassezia* species interactions to bacterial genera were plotted using ‘igraph’^27^ when p<0.05, as previously described^28^.

**Statistical analyses**

Alpha-diversity comparisons were performed on read-normalized, species-level data using ‘vegan’^29^. Other comparisons were performed on strain-, species-, or genus-level data, as indicated for each analysis. Fisher’s Exact tests^30^ were performed with a custom Python script. Wilcoxon test was performed with ‘stats’^31^ in R or Prism software (GraphPad Software, Boston, MA, USA). LEfSe^5^ (v1.1.1) was used to determine significant linear relationships of fungal or bacterial genera or species between Yanomami and Western data (HMP or Western expeditioner, as indicated in each analysis). We used a 50% taxa prevalence cutoff to filter bacterial and fungal taxa for comparisons between the Expeditioner and Yanomami across all body sites. Because ages are approximated for Yanomami participants, we assigned participants to age groups when evaluating age-related effects. Age groups reflect both biological (chronological) age and community roles/responsibilities, which may influence daily activities. Dunn tests^32^ (‘dunn.test’) with Benjamini-Hochberg (B-H) correction were used for multiple comparisons following Friedman test (for repeated measures, non-parametric analysis of variance (ANOVA)) group-wise comparisons. Paired body site comparisons of groups comprising >1 body site (e.g., combined sebaceous sites) or comparisons among sample groups comprising multiple time points (e.g., expeditioner’s pre-Amazon, in-transit, or post-Amazon samples during the 2018 expedition) were performed using ANOVA of linear mixed effects modeling (LME^33^; ‘lmerTest’), followed by Tukey’s method for multiple comparisons^34^ using ‘multcomp’. Multivariate data comparisons (e.g., taxa relative abundance and functional relative abundance) were performed using permutational analysis of variance (PERMANOVA) on Bray-Curtis (BC) dissimilarity matrices (square root-transformed taxonomic data) and visualized using Principal Coordinates Analysis (PCoA). Nested PERMANOVA (subject nested within age group: BC dissimilarity ~ Body Site + Age group + Expedition + Subject) used to evaluate bacterial or fungal community composition among all Yanomami samples (n=94) was performed using PERMANOVA+ software (PRIMER-e, Auckland, New Zealand). Heatmaps^35^ (‘gplots’) were generated using prominent taxa (bacteria: 80% prevalence; fungi: 50% prevalence), and distribution differences between Yanomami and HMP RAC samples were tested using Fisher’s Exact test followed by false discovery rate (FDR) adjustment for multiple comparisons (custom python script).

<https://CRAN.R-project.org/package=Hmisc>.
